# Hippocampal representations of partner and novel individuals in monogamous California mice during pair bond formation

**DOI:** 10.64898/2026.05.08.723922

**Authors:** Kimberly Hernandez-Palacios, Oshmita Golam, Steven A. Siegelbaum, Andres Bendesky

## Abstract

The hippocampal CA2 region is critical for social novelty recognition memory—the discrimination of whether a conspecific is novel or familiar. However, its role in forming a memory of a pair-bonded mate is unknown. To examine how social memories of pair-bonded individuals are encoded, we sought to understand if CA2 and the neighboring CA1 region participate in the memorization and recognition of a pair-bonded mate in monogamous *Peromyscus californicus* (California mice). Here, we report that CA2 and CA1 show distinct changes in social encoding of an opposite sex conspecific following pair-bonding. Using multi-channel silicon probes, we recorded single units from CA2 and CA1 in freely behaving male mice before and after pair bond formation during interactions with novel and partner females. We found that the strength of CA2 representations of a novel female mouse weakened after pair bond formation, indicating that CA2 may be preferentially important for novelty detection. In contrast, CA1 demonstrated an increase in the strength of encoding a female partner after pair-bond formation, suggesting that CA1 may encode partner memory. These findings indicate that pair bonding shifts the discrimination of social information from CA2 to CA1.

## INTRODUCTION

For many species, the formation of social bonds is essential for survival, as they provide cohesion useful for cooperation and collective success. The pair-bond between sexual partners in monogamous species presents a unique model to understand the neural circuits involved in social bonds. This exclusive bond becomes evident by the behaviors pair-bonded animals display towards their partner compared to behaviors towards other conspecifics. For example, pair-bonded animals huddle peacefully with their partner, fight novel individuals, and provide joint care for their offspring^1–4^. Like other monogamous mammals, monogamous species of *Peromyscus*, including *P. polionotus* (oldfield mouse) and *P. californicus* (California mouse), exhibit robust pair-bonding behaviors^5–9^. Notably, the California mouse shows, by many behavioral metrics, the most pronounced pair-bonding behavior of any mammalian species^10, 11^.

Several brain regions supporting the formation of pair-bonds have been identified, such as the nucleus accumbens^12^ and hypothalamus^13^. In addition, multiple lines of evidence implicate the neuropeptides oxytocin^13–15^ and vasopressin^16^. Although the brain regions involved in encoding the memory of a partner have not been identified, the hippocampus is known to be important for the encoding of different forms of social memory, including the discrimination between familiar and novel individuals. Two hippocampal areas are particularly important for social novelty recognition memory: the CA1 region in the ventral portion of the hippocampus^17^ and the CA2 region in the dorsal portion^18^, as well as the connections from dorsal CA2 to ventral CA1^19^. Silencing dorsal CA2^18, 20, 21^, ventral CA1^17^, or the projections from dorsal CA2 to ventral CA1^19^, inhibits the ability of a mouse to discriminate between a novel and a familiar conspecific. CA2 also regulates the innate behavior of social aggression by providing a social novelty signal that enhances aggression to a novel intruder compared to a familiar intruder^27^. Electrophysiological and calcium imaging studies have identified populations of neurons in ventral CA1^17^ and dorsal CA2^22–25^ that encode mouse identity and discriminate a novel from a familiar individual, consistent with the role of these regions in controlling social memory behaviors. The role of dorsal CA1 is somewhat more controversial, with some studies suggesting it is not required for social discrimination^17, 22^ and others indicating its importance in associating a conspecific with reward^26^. However, little is known as to whether and how the hippocampus encodes the memory of a partner.

That CA2 might help the formation of a pair bond is suggested by the findings that CA2 receives input from vasopressin and oxytocin neurons in the paraventricular nucleus of the hypothalamus and sends output to the lateral septum^28^, two regions known to be important for pair bonding^29, 30^. Moreover, activation of the vasopressin input to CA2 enhances both social memory and social aggression^31^. Finally, chemogenetic activation of CA2 promotes pair-bond-like behaviors in the promiscuous house mouse, *Mus musculus*^32^. However, whether CA2 encodes pair-bond memories in species that normally form pair bonds is unknown. To examine this question, we characterized dorsal CA1 and CA2 activity from male California mice as they form pair bonds with female partners.

## RESULTS

### Measuring pair-bonding behaviors in California mice

To measure pair-bonding behaviors in California mice (**Figure 1a**), we employed a partner-preference test modified from one commonly used in prairie voles^13, 33, 34^. In this assay (**Figure 1b**), a subject explores a partner and a novel opposite-sex mouse tethered to opposite corners of an open arena for 60 min (**Figure 1b**). Before the test, pairs were co-housed for a week, during which time they mated and had sufficient time for pair-bond establishment^35, 36^. We then measured selective affiliative contact towards the partner and aggression towards a stranger during the partner preference test (**Figure 1c**)—signature behaviors of pair-bonded California mice^5–8^.

**Figure 1.**
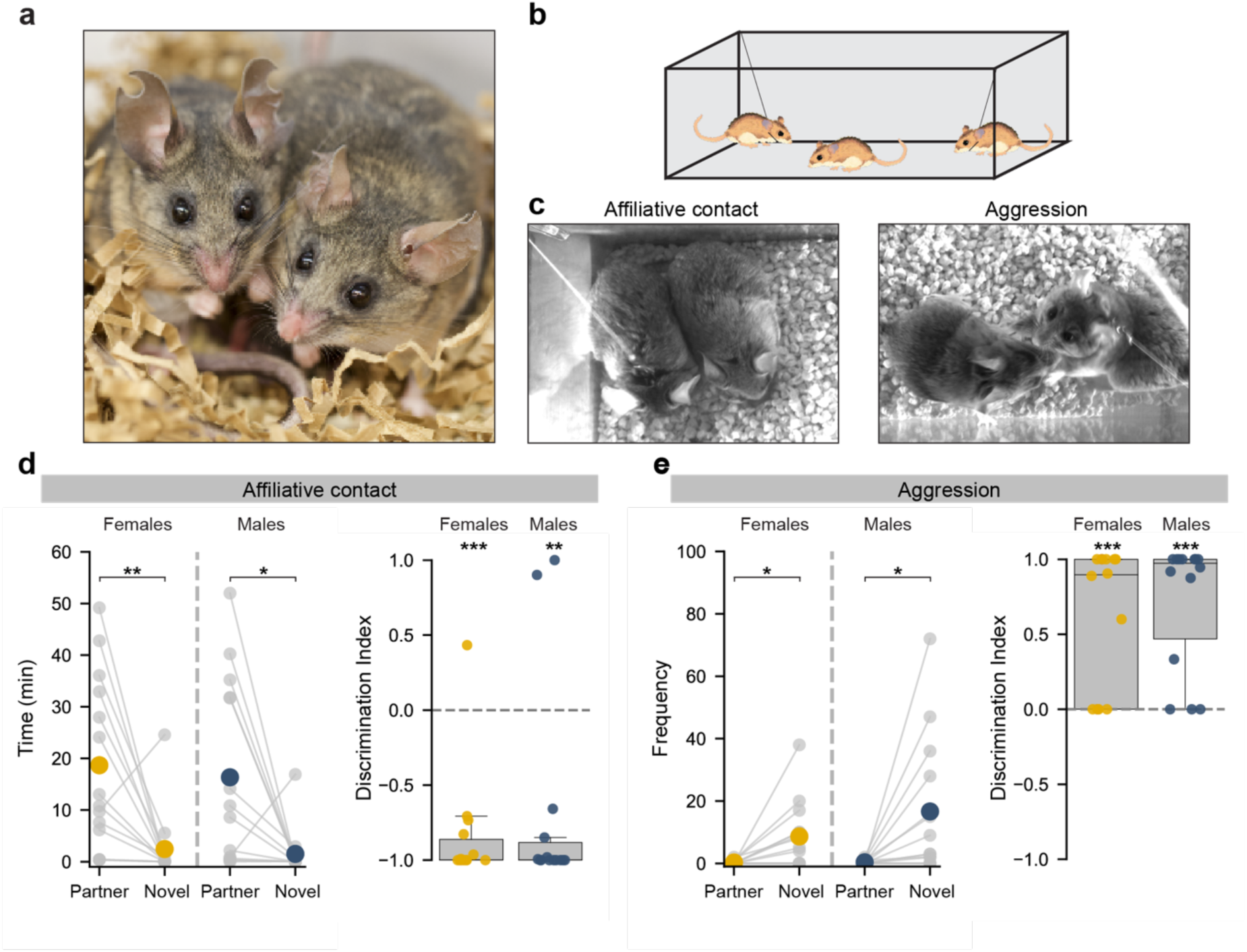
California mice display robust pair-bond behavior. **a)** Pair-bonded *Peromyscus californicus* (California mice) pair. **b)** Tether pair-bond assay. A male or female test mouse is allowed to interact with a tethered opposite-sex partner or a novel opposite-sex conspecific after pair-bonding. **c)** Pair-bond measured through differences in affiliative contact and aggression towards a partner versus novel mouse in a 60 min test session. **d)** (Left) Both female and male California mice display increased total affiliative contact time during a session with the partner compared to novel mouse. Females: paired t-test, t(13) = 3.308, *p* = 0.0057, n=14; Males: paired t-test, t(13) = 2.820, *p* = 0.0145, n= 14. (Right) Affiliation discrimination index for partner versus novel mouse interaction times. Discrimination Index = (novel - partner)/(novel + partner) affiliative contact times during session. Females: One-sample t-test against 0, t(13) = −8.258, *p* = 1.5771e-06, n=14; Males: One-sample t-test against 0, t(13) = −3.661, *p* = 0.0029, n = 14. **e)** (Left) Females and males show increased number of aggression bouts during session with novel stimulus mouse compared to partner. Females: paired t-test, t(13) = −2.970, *p* = 0.0109, n=14; Males: paired t-test, t(13) = −2.822, *p* = 0.0144, n= 14. (Right) Aggression discrimination index for novel versus partner. Discrimination index = (novel – partner)/(novel + partner) number of aggression bouts per session. Females: One-sample t-test against 0, t(13) = 4.720, *p* = 0.0004, n=14; Males: One-sample t-test against 0, t(13) = 6.308, *p* = 2.7115e-5, n = 14. Boxplots indicate median and interquartile range. \**p* < 0.05, \*\**p* < 0.01, \*\*\**p* < 0.001.

Both female and male California mice displayed 9-fold more affiliative contact towards a partner than to an opposite-sex novel mouse (females: paired t-test, *p* = 0.0057, n=14; males: paired t-test, *p* = 0.0145, n= 14; **Figure 1d**). Females and males also exhibited 30-fold more aggression towards a novel individual (females: paired t-test, *p* = 0.0109, n=14; males: paired t-test, *p* = 0.0144, n= 14) (**Figure 1e**). These results confirm the formation of pair-bonds under the conditions of our experiments, supporting the use of California mice as a suitable model for investigating the neural encoding of a pair-bonded partner.

### Hippocampal single unit recordings during pair-bond formation

Prior to recording neural activity from CA1 and CA2, we first asked whether a molecularly defined CA2 region existed in California mice, using well-defined CA2 markers in house mice: Purkinje cell protein 4 (PCP4) and striatum-enriched protein-tyrosine phosphatase (STEP) ^18, 38, 39^. Using immunohistochemistry, we found that these markers were colocalized in a relatively small hippocampal region wedged between CA1 and CA3, similar to CA2 in other species (**Figure 2a**). Using these markers, we defined stereotaxic coordinates to target four-shank 64-channel silicon probe electrodes spanning the CA1 and CA2 regions (**Figure 2b, c**). We recorded neural activity from CA1, CA2, or both regions, depending on device targeting and verified probe localization postmortem.

**Figure 2.**
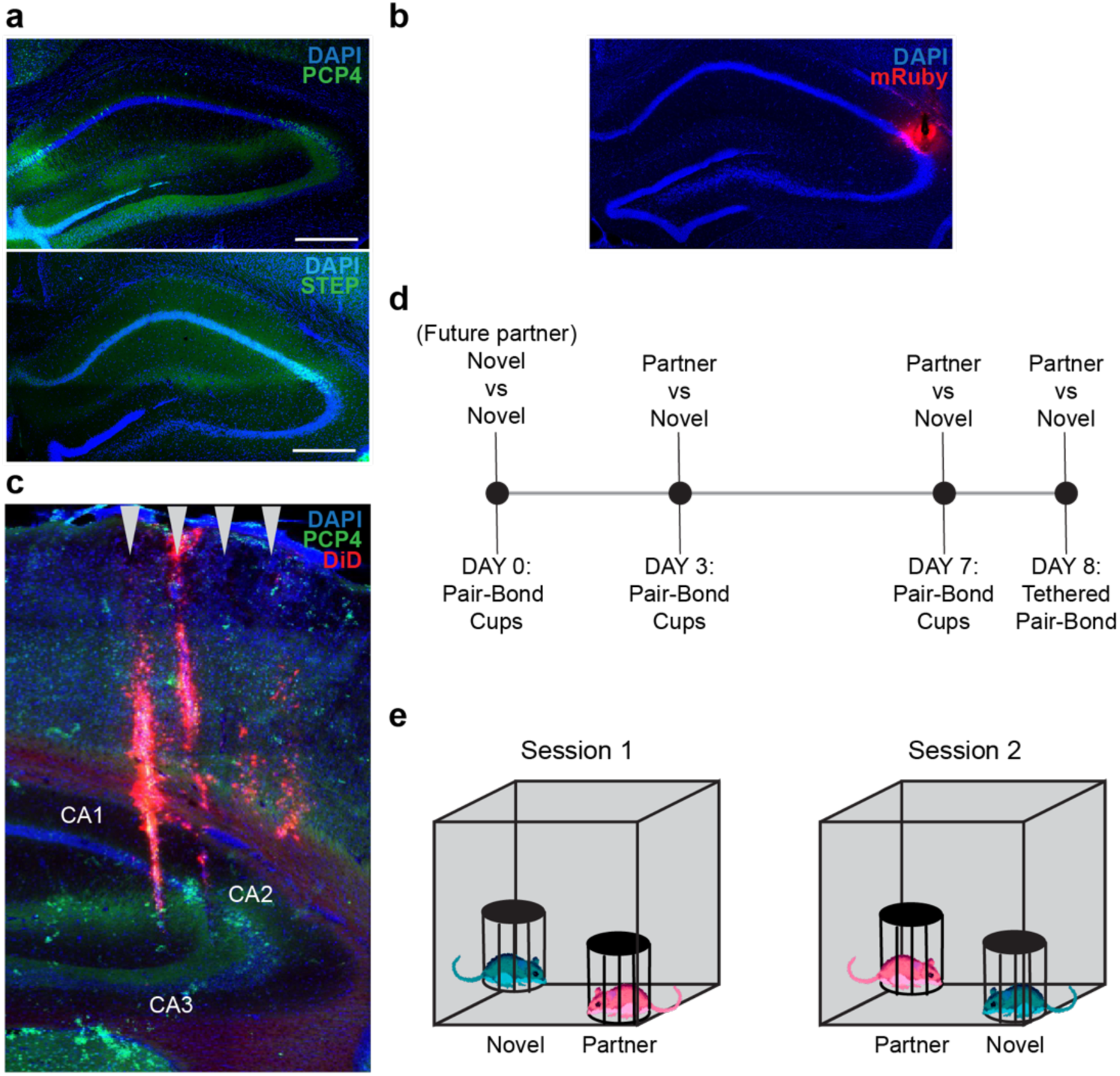
Experimental design for electrophysiological recordings. **a)** Confocal images of expression of CA2 markers PCP4 and STEP in California mice. Top: PCP4. Bottom: STEP. Scale bars: 500 µm. **b)** Targeted mini-Ruby (red) injections to confirm dCA2 coordinates in California mice. **c)** Post hoc assessment of targeting of silicone probes into the CA1/CA2 region of the hippocampus. DiD (red) was applied to identify probe location and co-stained with CA2 marker PCP4 (green). Red tracks show location of the four silicon probes in a representative recording. **d)** Recording timeline. On day 0, the subject male investigates two novel females. Future-partner was selected at random prior to testing. Partner mice cohabitate with subject mouse for 8 days after day 0 recording. Additional recordings were collected on days 3 and 7. On day 8, pair-bond strength was assessed in tethered pair-bond task. **e)** Pair-bond cups (PBC) assay. Neural activity was recorded as the subject explored two female mice confined to wire cup cages in two 10-min sessions. In the second session, positions of the stimulus mice were swapped.

To examine how CA1 and CA2 in males represent female individuals and how these representations are altered by pair bonding, we collected single unit extracellular electrophysiological data at three time points during the process of pair bond formation: days 0, 3 and 7 (**Figure 2d**). On day 0, prior to pair bond formation, a male mouse freely investigated two novel females, with one designated at random as the “future-partner” and the other as “novel”. This future-partner female was then co-housed with the subject male mouse in its home cage for the remainder of the study. We video recorded mice continuously to identify day of mating (**Figure S1a**). By day 3, 8 out of 13 pairs had mated. By day 7, 10 out of 12 pairs mated (one test mouse was removed due to damage to implant prior to day 7).

Because animals spent more time with their partner than with a novel individual in the tethered pair-bond assay (**Figure 1d, e**), the amount of interaction time and, thus, neuronal data that can be obtained with the novel mouse was limited. To facilitate comparisons of the neural representations of the partner and novel animal, we recorded neural activity as the male subject explored its partner and the novel mouse, each confined in wire cup cages placed at opposite corners of a square arena, as previously described for house mice^23, 24^. This allows the subject to freely explore both mice but prevents huddling or aggression, resulting in equivalent interaction times with partner and novel mice (**Figure S1b-g**). On each day, we recorded neural activity in two 10-min sessions, with the positions of the stimulus mice reversed in each session to disentangle spatial from social information encoding^23, 24^ (**Figure 2e**).

To confirm pair-bond formation in the implanted mice, we conducted the tethered pair-bond assay on the eighth day. All mated pairs (mated: n=10 out of 12) showed preferential affiliative contact with the partner over a novel female (paired t-test, *p* = 0.02; **Figure S2a**). One pair of non-mated (NM) mice showed preferential affiliative contact with one another whereas the second pair did not. Mated mice also engaged in selective aggression towards the novel female (paired t-test, *p* = 0.0510**; Figure S2b**).

### CA1 and CA2 neuron responses to novel and partner mice

We first examined mean firing rate responses for the population of recorded neurons during interactions with the partner (or future-partner) compared to the novel mouse, for both CA1 and CA2 before (day 0) and during the process of (days 3 and 7) pair bond formation (**Figure S4**). We classified cells as either excitatory or inhibitory based on waveform shape and autocorrelograms of the units (**Figure S3**). Because the number of inhibitory neurons was limited, most of our analyses focus on firing of excitatory neurons.

A previous study found a higher mean firing rate for CA2 neurons in male house mice during interactions with a novel same-sex animal compared to a familiar littermate^22^. Accordingly, we first compared mean firing rates averaged over all excitatory neurons during periods of interaction with a novel female compared to the partner. Across days, CA1 demonstrated comparable mean firing rates to both stimuli (**Figure S4b**). However, CA2 neurons in California mice showed a significantly higher firing rate around the novel mouse (3.05 Hz) compared to the partner (2.71 Hz) when assayed after pair bonding on day 7 (paired t-test with Bonferroni correction, p=0.0068). In contrast, no significant difference in CA2 firing rates were observed on day 3, suggesting that a week of co-housing may be necessary to establish strong familiarity encoding. Additionally, the variable time of mating allowed us to separate mice in different bonding states and examine the effects on CA2 activity. On day 3, CA2 neurons of mated males showed higher overall firing rates during interactions with either novel or partner stimuli compared to firing rates of unmated males (Welch’s t-test, Novel: *p* = 10^-7^, Partner: *p* = 10^-8^). This suggests that mating increases CA2 activity in response to conspecifics.

Next, we asked how individual CA1 and CA2 excitatory neurons respond during exploration of the two stimulus mice across the days of the experiments. To that end, we aligned the responses of CA1 and CA2 neurons to the start of each interaction with the stimulus mouse (**Figure 3a, b**). For each neuron, we measured its mean z-scored firing rate in 250 ms bins during a window starting five seconds before the onset of each interaction and ending five seconds after the onset of each interaction. We then measured the response of each cell during interaction with a given stimulus mouse based on the difference between the cell’s mean firing rate averaged 0 to 5 s after the start of the interaction with the mean firing rate averaged 5 to 1 s before the start of the interaction. When firing rates for all cells were displayed in raster plots ranked by the extent to which a given cell’s firing rate was increased during exploration of the partner (or future partner) on a given day, a fraction of cells showed a clear increase in activity during the interaction while the activity of other cells was inhibited. This pattern was less apparent when we plotted responses of the same neurons during interactions with the novel mouse in the chamber, suggesting that some neurons may be mouse-selective (**Figure 3a,b**; **Figure S5-S8**). Conversely, when we ranked neurons based on the strength of activation around the novel mouse in the chamber, we also saw populations of neurons that appeared to selectively increase their activity around this mouse and not around the other mouse.

**Figure 3.**
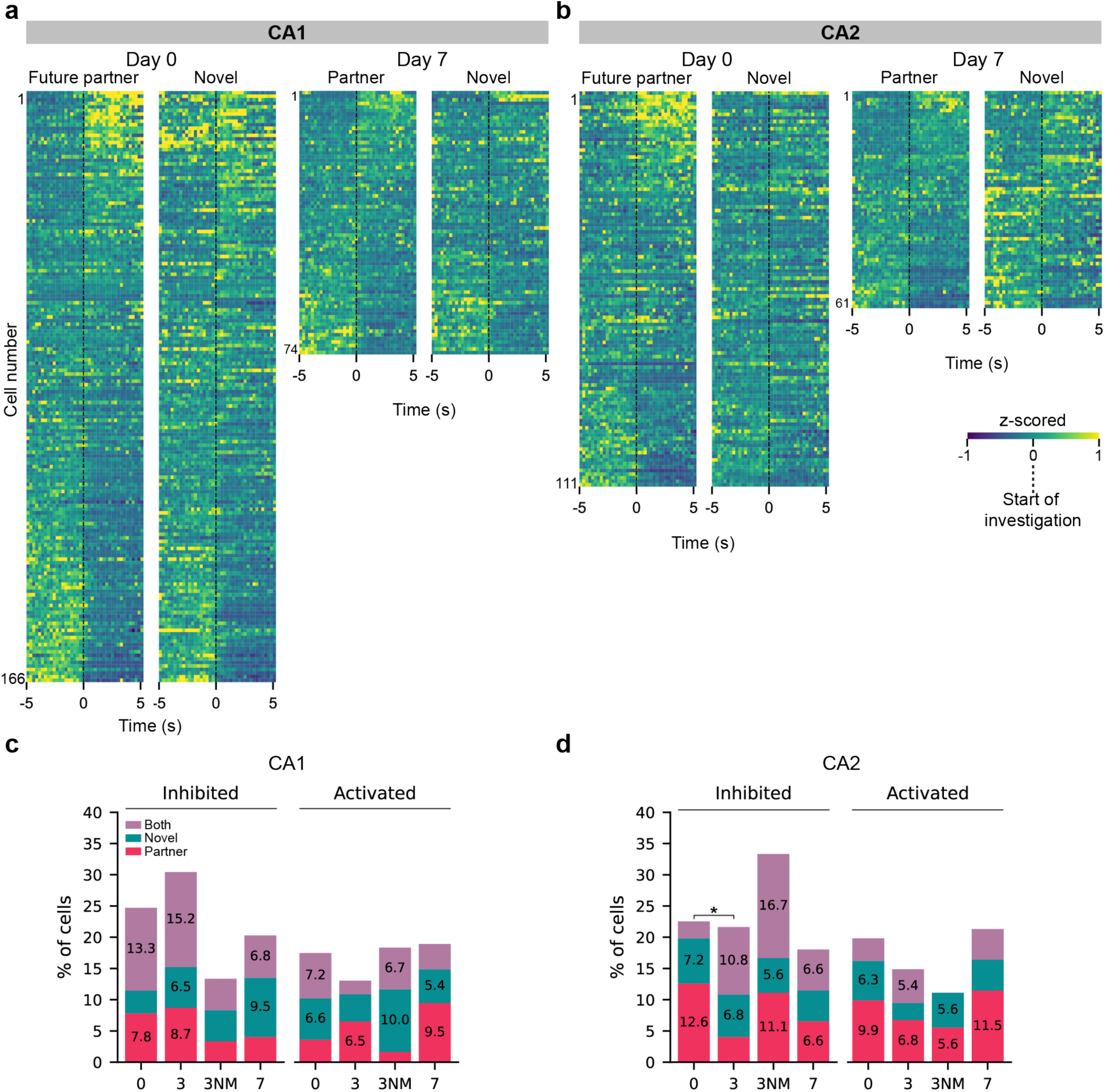
Cell firing responses to partner and novel mice. **a,b)** CA1 (a) and CA2 (b) excitatory neuron responses to stimulus mouse investigation on day 0 (left), before pair-bonding, and day 7 (right), after pair-bonding. Z-scored firing rates aligned to the onset of stimulus investigation (time 0). Each row shows the color-coded Z-scored activity of a neuron from indicated region, starting 5 seconds before onset of investigation to 5 seconds after. Cells are ranked by response strength to future-partner (day 0) or partner (day 7). Ranking was determined by the change in firing rate (FR) during social interaction relative to baseline, calculated as (FR_after – FR_before), averaging over windows t= 5 to 1 s before interaction and t= 0 to 5 s after start of interaction. (Day 0: CA1, n = 7 mice, 166 cells; CA2, n = 8 mice, 111 cells. Day 7: CA1, n = 4 mice, 74 cells; CA2, n = 4 mice, 61 cells). **c, d)** CA1 (c) and CA2 (d). Percent of neurons whose firing rate was significantly inhibited (decrease >97.5% of null distribution) or activated (increase >97.5% of null) during social interactions, relative to baseline, with partner only (pink), novel mouse only (teal) or with both (purple). Significance assessed by comparison of observed firing rate difference pre- and post-onset of investigation of a stimulus mouse with null distribution generated by pre-/post-onset bout label shuffling. Day-to day comparisons of response distributions: *, *p* <0.05, two-sided Fisher’s exact test, 2×4 table.

To provide quantitative insight into these cellular responses, we performed a statistical permutation test to identify neurons whose activity during interactions with a given stimulus mouse was significantly increased or decreased relative to baseline. For each neuron we further classified whether it was activated or inhibited selectively during interactions with one of the two mice, or both (**Figure 3c,d**). Across the days of the experiment, a fraction of neurons in CA1 and CA2 (∼10-15%) were significantly and selectively activated around either the partner or the novel mouse; around 5-7% of neurons were activated non-selectively around both stimulus mice. A somewhat greater fraction of neurons was inhibited during the social interactions, with around 10-20% showing selective inhibition and 5-15% showing non-selective inhibition. When we examined how responses changed over the days of the experiment, we observed a statistically significant change in category distribution of inhibited CA2 cells from day 0 to day 3 (Fisher’s exact test, *p* = 0.0371). When comparing CA1 and CA2 responses, the only statistically significant differences was that CA1 had a greater fraction of cells that were inhibited by both stimuli (13.3%) than did CA2 (2.7%) on day 0 (Fisher’s exact test, *p* = 0.0060).

### Comparison and dynamics of CA1 and CA2 responses to novel and partner mice

The above results provide a binary measure of response selectivity but do not incorporate differences in the magnitude of the firing rate changes during interactions with the two stimulus mice. To examine firing rate response selectivity, we measured the difference in mean firing rate changes around the two stimulus mice (firing rate around partner or future partner minus that around the novel mouse). We assessed the statistical significance of this difference by comparing it to a null difference distribution generated by random shuffling stimulus mouse labels 1,000 times. We classified a neuron as partner-selective if its observed firing-rate difference fell in the upper 2.5% of the null distribution (i.e., greater firing during partner investigation) and novel-selective if it’s difference fell in the lower 2.5% of the distribution. Cells in the middle 95% of the distribution were labeled as non-selective. We balanced the number of bouts per stimulus type within each session to achieve equivalent representation for each neuron.

We observed a striking change in the fraction of CA1 and CA2 cells selective for the novel mouse or partner (or future partner) across the 7 days of the recordings, with cells in the two regions showing opposing changes (**Figure 4a**). For CA1, the fraction of partner-selective cells increased markedly from 3% of cells on day 0 to 16.2% of cells on day 7. By contrast, the fraction of novel-mouse-selective cells showed the opposite effect, decreasing from 9.6% on day 0 to 4.1% on day 7. As a result, the ratio of partner-to-novel selective cells increased 13-fold, from 0.3 on day 0 to 4 on day 7.

**Figure 4.**
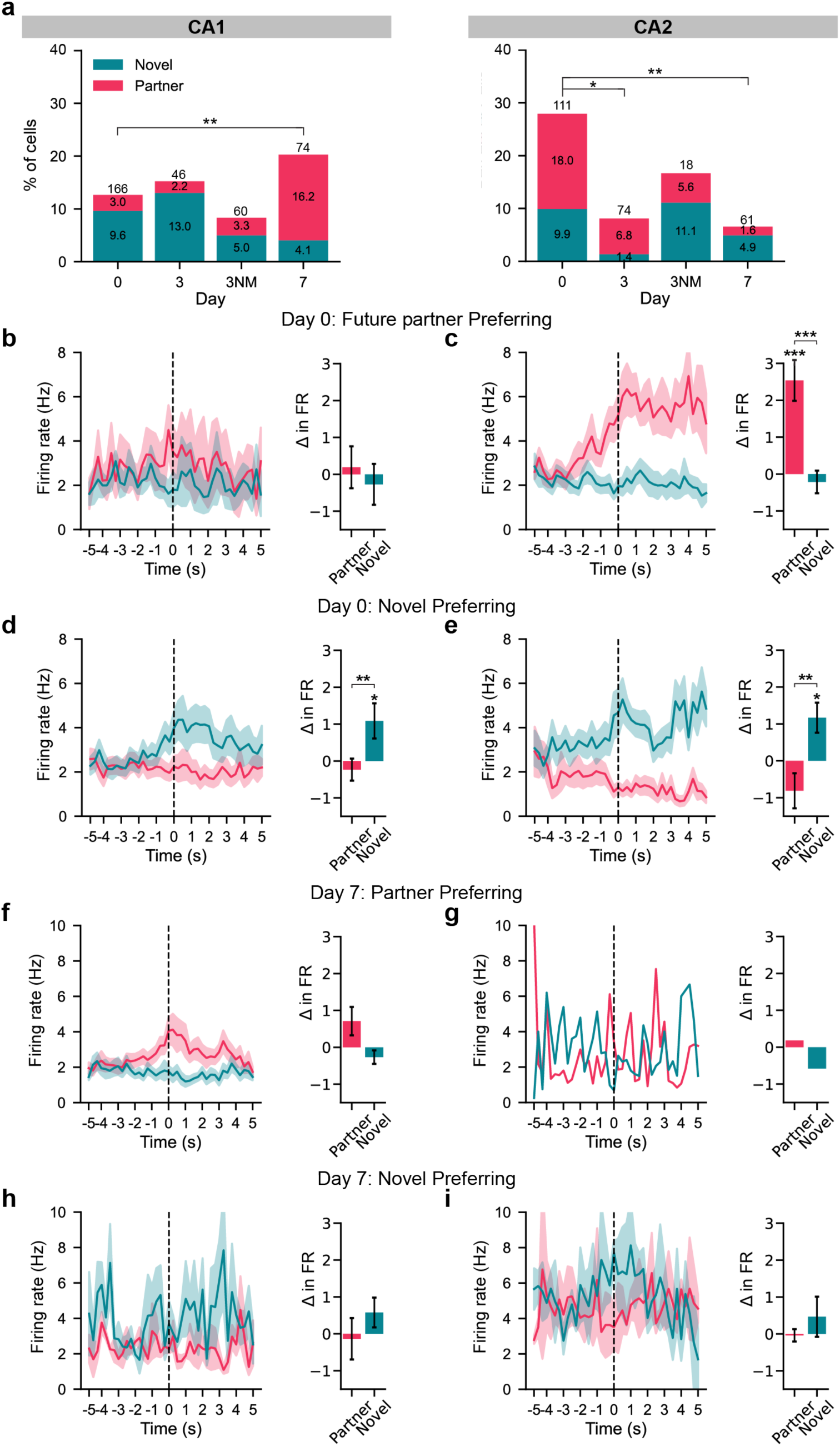
Stimulus-selective responses during partner and novel investigations. **a)** Percent of novel- and partner-preferring CA1 (left) and CA2 (right) excitatory cells on indicated recording days. Day 3 recordings grouped according to whether the pair had or had not mated (NM). Novel and partner-preferring cells defined by whether (partner FR - novel FR) difference was, respectively, ≥ 0.975 or ≤ 0.025 of shuffled null difference distribution. Percent cells with given preference shown inside bars. Significant effect of day on CA1 and CA2 preference distribution. CA1: Two-sided Fisher’s exact test, 3×4 table, *p* = 0.0041. Post hoc pairwise comparisons (Bonferroni-corrected) revealed significant differences between Day 0 and Day 7 (*p* = 0.0045). (n = 9, 4, 5, 5 mice for days 0, 3, 3NM, and 7 respectively). CA2: Two-sided Fisher’s exact test, 3×4 table, *p* = 0.0008. Post hoc pairwise comparisons (Bonferroni-corrected) revealed significant differences between Day 1 and Day 3 (*p* = 0.0153) and between Day 1 and Day 7 (*p* = 0.0054). (n = 8, 5, 3, 4 mice for days 0, 3, 3NM, and 7 respectively). **b-i)** Left panels, Mean FR during interactions with partner (or future partner) (pink) or novel mouse (teal) from day 0 and 7 stimulus-selective cells. Mean FR aligned to start of interaction (dotted line). Shaded region: SEM. Right panels, FR change during interaction relative to −5 to −1 s baseline average. **b,c)** Responses on day 0 for CA1 (b) and CA2 (c) future-partner-selective cells during interactions with future partner and novel mouse. CA2 FRΔ: Paired t-test between partner and novel, t(19) = 4.832, *p* = 0.0001; n=20 cells. One-sample t-test against zero: partner, t(19) =4.591, p = 0.0002. CA1 FRΔ: ns. **d,e)** Responses on day 0 for CA1 (d) and CA2 (e) novel-mouse-preferring cells during interactions with future partner and novel mouse CA1 FRΔ: Paired t-test between partner and novel, t(15) = −4.210, *p* = 0.0008; n=16 cells. One-sample t-test against zero: Novel, t(15) = 2.286, p = 0.0372. CA2 FRΔ: Paired t-test between partner and novel, t(10) = −4.796, *p* = 0.0007; n=11 cells. One-sample t-test against zero: Novel, t(10) = 2.861, p = 0.0169. **f,g)** Mean responses on day 7 for CA1 (f) and CA2 (g) partner-preferring cells during interactions with partner and novel mouse (ns for both). **h,i)** Mean responses for novel preferring cells on day 7. n=3 cells for CA1(h) and CA2 (i) (ns for both). Error bars represent ± SEM. \**p* < 0.05, \*\**p* < 0.01, \*\*\**p* < 0.001

In contrast to CA1, CA2 showed a decrease in the fraction of both partner-selective and novel-mouse-selective cells across the 7 days of the experiment. The fraction of partner-selective cells decreased 11-fold, from 18% on day 0 to only 1.6% on day 7. The fraction of novel-mouse-selective cells also decreased, although less dramatically, by only 2-fold, from 9.9% on day 0 to 4.9% on day 7.

Comparison of stimulus preference between CA1 and CA2 revealed a significant difference in distributions on day 0 (Fisher’s Exact test, *p* = 10^-5^). A greater fraction of partner-preferring cells was present in CA2 than CA1 (Fisher’s Exact test, Bonferroni-corrected, *p* = 10^-5^). Cell preference also differed on day 7 between the regions (Fisher’s exact test, *p* = 0.0092), with an enrichment of partner-preferring cells for CA1 compared to CA2 (Fisher’s exact test Bonferroni-corrected, *p* = 0.0186). These results demonstrate a shift in partner-preferring cells from CA2 to CA1 after pair-bond formation.

The unequal number of future-partner-selective and novel-mouse selective cells on day 0 in both CA1 and CA2 was unexpected. As the two groups of females were equally novel to the male and assigned to the future partner group at random, these differences likely reflect some random variation in the sensory cues of the females in the two groups. For example, it might be related to a compatibility preference that has been identified upon initial exposure to potential mates in *Peromyscus*^40, 41^. Another possibility we considered is that the mice in the two groups differed in the distribution of estrus cycle stage. However, there was no correlation between estrus cycle and neural social information encoding.

To obtain a more quantitative readout of response magnitude and dynamics, we measured the time course of mean firing rates before and during the social interactions for the population of partner-selective and novel-mouse-selective neurons in both CA1 and CA2. We constructed separate event-triggered averages for the two classes of neurons aligned to the start of mouse interactions, spanning 5 sec before and 5 sec after, across experimental days (**Figure 4b-i**).

On day 0 the sparse population of future-partner selective CA1 cells showed a small firing rate response during interactions with the novel mouse that was not significantly different from baseline or from the firing rate response during interactions with the future partner (**Figure 4b**). In contrast, the population of novel-mouse selective CA1 cells showed a marked increase in average firing rate over baseline levels during interactions with the novel mouse, with no change in rate with the partner (paired t-test, *p* = 0.0008) (**Figure 4d**).

By day 7, we observed an increase in mean firing rate of the population of CA1 partner-selective cells during interaction with the partner but not the novel mouse; however, the change in firing rate from baseline was not significant (**Figure 4f**). The novel-mouse-selective cells showed no significant increase in firing around either the novel or partner mouse (Wilcoxon signed-rank test, *p* = 0.25; **Figure 4h**), consistent with the decrease in the fraction of novel-mouse-selective cells on this day (**Figure 4a**).

Co-housing produced an opposite change in the CA2 population firing rate responses compared to CA1, consistent with the distinct changes in the fraction of mouse-selective CA1 and CA2 neurons during this period (**Figure 4a**). On day 0, the population of CA2 future-partner-selective cells showed a large increase in mean firing rate relative to baseline during interactions with the future partner that was significantly greater than that seen during interactions with the novel mouse (paired t-test, *p* = 0.0001; **Figure 4c**). Conversely, the novel-mouse-selective cells showed a significantly greater increase in firing compared to baseline during interactions with the novel compared to partner mice (paired t-test, *p* = 0.0007; **Figure 4e**). These results are consistent with findings on house mice, where CA2 activity has been found to have distinct representations of two novel mice^24^.

By day 7, after pair-bonding, CA2 cells no longer increased their firing rate during interactions with either the partner or the novel mouse (**Figure 4g, i**). These results are consistent with the decrease in the fraction of CA2 mouse-selective cells during co-housing (**Figure 4a**). Together these findings indicate regional changes in both the fraction of stimulus-selective cells and their firing rate modulation around conspecifics following pair-bond formation.

### Decoding of mouse identity and location by CA1 and CA2 population activity

The above single-cell analyses relied on classifying information-coding properties of CA1 and CA2 based on changes relative to statistical thresholds. However, this approach will fail to detect contributions to information encoding by weakly-selective or mixed-selectivity neurons at the population level^42^. To determine whether CA1 and CA2 population activity encodes pair-bond information, we trained a support vector machine classifier with a linear kernel to decode interactions with the partner (or future-partner) compared to a novel conspecific before and after pair bond formation.

To achieve sufficient power and to balance neural contributions for cross-day comparisons, we trained the decoders on a pseudo-population of neurons from recordings from all mice, using the same number of neurons each day (based on the minimal number of neurons recorded on a given day and region). This resulted in our training the classifier using the activity from 46 CA1 and 61 CA2 neurons. Mean firing rates were measured in 250 ms time bins throughout the interactions with either stimulus mouse. To decode identity, we balanced location information by combining recordings around a given mouse across the two cup locations in the two sessions, ensuring that we included equal times of exploration around each cup. To decode location, we balanced mouse identity information by combining data during exploration around a given cup (left or right) from the two sessions, ensuring that we included equal amounts of time spent exploring the two mice in a given cup.

Decoding performance based on CA1 pseudo-population activity for both mouse identity and cup location was significantly greater than chance across all days (**Figure 5a**). Decoding accuracy was relatively unchanged across days, with location decoding tending to be more accurate than identity decoding. Thus, the encoding of mouse identity and location was stable in CA1, with no significant change during pair bond formation.

**Figure 5.**
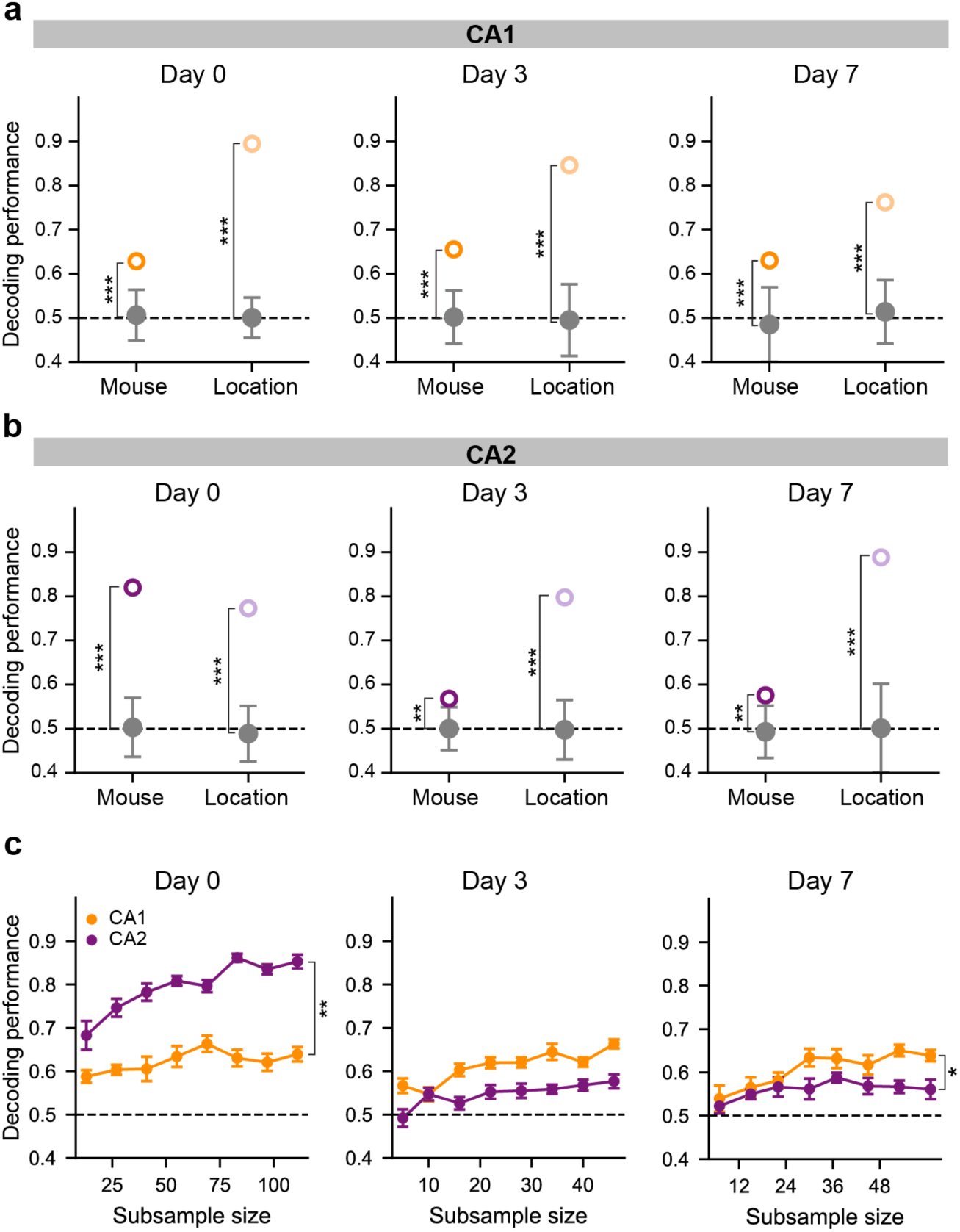
Effect of partner co-housing on mouse identity and location decoding accuracy by CA1 and CA2 pseudo-populations. Open symbols, accuracy of decoding mouse identity (darker circle) and location (lighter circle). Grey, null model decoding accuracy. **a)** Decoding of mouse identity and location based on CA1 activity. Day 0 mouse decoding = 0.63 vs. null model = 0.51±0.03; location decoding = 0.89 vs. null model = 0.50±0.02. (Pseudo-population of 46 cells, n = 9). Day 3: mouse decoding = 0.66 vs. null model = 0.50±0.03; location decoding = 0.85 vs. null model = 0.50±0.04. (Pseudo-population of 46 cells, n = 4). Day 7: mouse decoding= 0.63 vs. null model = 0.48±0.04; location decoding = 0.76 vs. null model = 0.51±0.04. (Pseudo-population of 46 cells; n = 5). **b)** Decoding based on CA2 activity. Day 0: mouse decoding = 0.82 vs. null model = 0.50±0.03; location decoding = 0.77 vs. null model = 0.49±0.03. (Pseudo-population of 61 cells, n = 8). Day 3: mouse decoding = 0.57 vs. null model = 0.50±0.02; location decoding = 0.80 vs. null model = 0.50±0.03. (Pseudo-population of 61 cells, n = 5). Day 7: mouse decoding= 0.58, vs.null model = 0.49±0.03; location decoding = 0.89 vs. null model = 0.50±0.05. (Pseudo-population of 61 cells, n = 4). **c)** Comparison of decoding accuracy of mouse identity across days based on CA1 or CA2 activity as a function of pseudo-population subsample size. Day 0, Decoding performance increased with subsample size and was greater in CA2 compared to CA1 (based on linear model examining region x subsample value interaction; Type II ANOVA, (F1, 12) = 9.79, *p* = 0.0087). Day 3, No significance difference in CA1 compared to CA2 decoding accuracy. Day 7, Decoding accuracy increased with subsample size and was greater in CA1 than CA2 (interaction: Type II ANOVA, (F1, 12) = 6.33, *p* = 0.0271). Error bars represent ± SEM. \**p* < 0.05, \*\**p* < 0.01.

By contrast, identity decoding based on CA2 activity declining markedly in accuracy over the 7 days of the experiment (**Figure 5b**). These results are in line with the pronounced decrease in the fraction of partner-selective CA2 cells from day 0 to day 7 (**Figure 3b**). The accurate decoding of mouse identity on day 0 is also consistent with previous findings that CA2 in house mice can discriminate two novel mice^24^. In contrast to the change in identity decoding, the classifier based on CA2 activity showed a stable decoding of location across days. These results indicate that pair-bonding is linked to a weakened ability of CA2 to discriminate individuals but not location.

Given the observed differences in decoding performance, we sought to determine the extent to which this performance was driven by stimulus-selective neurons versus contributions from weakly or non-selective cells. Specifically, we examined whether decoding accuracy for distinguishing a partner from a novel mouse depended primarily on a subset of highly selective neurons, or whether population activity from more weakly tuned neurons contributed meaningfully to classification. To address these possibilities, we trained separate classifiers on CA1 and CA2 datasets after excluding neurons previously identified as stimulus-selective (**Figure S9**). As a control, we trained additional classifiers on pseudo-populations in which an equivalent number of randomly selected neurons were removed. Following this procedure, the resulting pseudo-populations consisted of 39 neurons in CA1 and 57 neurons in CA2. Removal of stimulus selective cells resulted in a decrease in performance scores for stimulus decoding across days in CA1 (**Figure S9a**). Still, even after the removal of stimulus selective cells, decoding performance was greater than chance on days 0 and 3, the latter for both mated and non-mated groups. However, the exclusion of stimulus selective cells on day 7 reduced the ability of the classifier to decode partner and novel stimulus mice above chance levels. In CA2, deletion of stimulus-selective cells also reduced decoding performance across days. This reduction was most prominent on day 0, with a drop in performance from 0.83 to 0.56, leading to a failure to decode identity above chance (**Figure S9b**). Decoding performance after removal of stimulus selective cells was also not greater than chance on day 3. However, decoding performance was still significant on day 7 after the exclusion of these cells, indicating that weakly selective cells encoded stimulus identity. The removal of stimulus-selective cells did not affect location decoding across days in either region.

Finally, we quantitatively compared decoding performance in CA1 and CA2 using subsampling^43^ to obtain classifiers containing the same number of neurons, assessing decoding performance across regions as a function of the number of neurons in different sets of subsamples (**Figure 5c**). On day 0, identity decoding performance in CA2 was consistently greater than that in CA1 across the different subsample sizes (p= 0.0087). However, on day 7 this pattern was reversed, with CA1 decoding accuracy greater than in CA2 (p=0.0271). In contrast, there was no difference in location decoding performance between CA1 and CA2 (**Figure S10**). These results form a consistent picture with our single cell results that pair bond formation exerts opposing effects to strengthen social representations in CA1 while weakening social representations in CA2.

## DISCUSSION

Here, we report that the CA1 and CA2 regions of the hippocampus from male California mice generate representations capable of discriminating two novel females as well as a pair-bonded female partner from a novel female conspecific. Longitudinal electrophysiological recordings reveal a progressive reorganization of stimulus representations across CA1 and CA2 during pair-bond maturation, with CA2 preferentially discriminating the identity of novel individuals on day 0, prior to pair bond formation, and CA1 strengthening its representation of the familiar partner on day 7 of co-housing, after formation of a pair-bond.

A potential role for the hippocampal CA2 region in mediating pair-bond formation had been suggested based on the finding that chemogenetic activation of CA2-projecting paraventricular nucleus neurons in house mice significantly increased side-by-side contact time with a familiar partner relative to a novel conspecific^32^. However, the significance of this finding to true pair bonding in monogamous species is unclear. Our findings in CA2 recordings from California mice suggest that rather than contributing to representations of pair-bonded animals, CA2 may serve to discriminate mouse identity prior to pair bonding, with CA1 forming the long-term partner representation.

Previous work in male house mice revealed that CA2 social representations could distinguish the identity of two male conspecifics, regardless of whether they were familiar or novel^24^. Here we find that CA2 of California mice encodes distinct representations of two novel opposite-sex conspecifics, as evidenced by the large number of stimulus-mouse-selective cells and high accuracy of decoding mouse identity using data from day 0 interactions, when both stimulus mice were novel. The future-partner and novel-mouse selective cells were highly responsive for their respective categorization, as their average firing rates were substantially increased at the onset of stimulus investigation. These findings are in broad agreement with the role of CA2 in social identity discrimination of novel animals in house mice^18, 20, 27, 44^.

Based on the established role of CA2 in discriminating familiar from novel conspecifics^18–20, 22, 24^ and its necessity for the recall of well-established social memories^18, 19^, we anticipated a significant *increase* in the proportion of partner-selective cells to that of novel-mouse-selective cells following pair-bonding, given that a pair-bonded partner is likely to possess greater salience than a novel individual.

Contrary to this expectation, CA2 exhibited a significant *decrease* in the number of partner-selective cells during the one week of co-housing. It is possible that CA2 becomes less responsive in identifying novel from familiar conspecifics after pair-bonding, as novel animals might lose salience. Given that our first recordings after the start of co-housing were performed on day 3, it would be interesting to examine how quickly this change develops over the 3 days.

Previous work in house mice also found that dorsal CA1 neural activity contained weaker representations of same-sex mouse identity compared to CA2^17^. Moreover, dorsal CA1 in house mice is not required for the behavioral discrimination of a novel versus familiar conspecific^17, 22^. We find similarly that dorsal CA1 population activity in California mice can accurately decode the social identities of two novel opposite-sex conspecifics on day 0, with a decoding accuracy lower than that seen in CA2 on this day. CA1 can also decode a novel compared to familiar pair-bonded conspecific on days 3 and 7, the latter showing a greater fraction of partner-selective cells. The activity from these partner-selective cells is required for the decoding of interactions with the partner versus the novel mouse, indicating that CA1 encodes social information partly through a subpopulation of mouse-selective cells. Our results on dorsal CA1 social encoding in California mice are reminiscent of findings for ventral CA1 in house mice, which is important for social identity recognition of both same^17^ and opposite-sex^45^ house mice. Whether the difference in dorsal CA1 social-response activity between house and California mice is species- or sex-related remains to be determined.

Through single cell analyses, we further uncovered how the responses to conspecifics evolve as the pair-bond forms. While the ratio of partner to novel selective cells remains relatively stable in CA1 from day 0 to 3, we observed a striking increase in partner selective cells on day 7. This suggests a potential role of CA1 in the recognition and representation of a pair-bonded partner. It is further consistent with the idea that the memory-based firing changes upon pair bond formation may take several days to develop after mating. The increase in partner-selective cells we observe following pair-bonding in dorsal CA1 is reminiscent of the increase in dorsal CA1 cells selective for a same-sex mouse associated with a water reward^26^. This suggests the possibility that the increase in representation of partner-selective cells in dorsal CA1 following mating reflects the association of the partner with a pair-bonding reward.

The above results indicate a shift in the discrimination of social information from CA2 to CA1 during pair-bond formation. Whereas CA2 has a high population decoding performance and a large number of stimulus selective cells when an animal first meets two individuals of the opposite sex, co-housing leads to a marked decrease in both measures of social information. In contrast, CA1 shows an increase in the fraction of social identity-selective cells over this time. As noted above, these results suggest that CA2 may be preferentially involved in the encoding of the identity of novel individuals and that CA1 may be more important for long-term memory storage of social identity of familiar individuals.

Our finding that population decoding performance in CA1 is unchanged by pair bonding despite the increased number of identity-selective cells suggests that the CA1 population code may switch from a reliance on weakly- or mixed-selective neurons prior to pair bond establishment (days 0 and 3) to a reliance on identity-selective cells after pair bonding (day 7). This idea is consistent with our finding that removal of the identity-selective CA1 neurons from the population decoder does not decrease decoding accuracy on days 0 and 3 but does lead to decoding failure on day 7. In contrast, decoding based on CA2 population activity relies more heavily on the identity-selective cells on days 0 and 3 compared to day 7.

Exactly how pair-bonding results in a shift in social information processing from CA2 to CA1 and to a reliance on identity-selective compared to weakly or mixed selective neurons remains to be determined. One possible mechanism may occur through alterations in expression of vasopressin and oxytocin receptors^30^, as these receptors have been implicated in social novelty recognition in CA2 and other brain regions^46^. The action of such neuropeptides may lead to distinct plastic changes in neural connectivity of CA1 and CA2 neurons, resulting in distinct shifts between highly-selective (low-dimensional) and mixed-selective (high dimensional) neural responses.

Although our study did not examine hippocampal projections in the California mouse, our findings point toward a potential memory mechanism that influences reward valuation. Previous work has demonstrated that dorsal CA2 projects to ventral CA1, which in turn sends outputs to the nucleus accumbens (NAc) shell^19^. Research in prairie voles has highlighted the critical role of mesolimbic reward circuits, including the NAc and ventral tegmental area (VTA), in mediating pair-bond formation and maintenance^12^, ^47–49^. Studies have also identified reward associated neurons in dorsal CA1^24, 50^. We suggest that hippocampal outputs, particularly through the dCA2-vCA1-NAc pathway, may interact with reward circuits to assign positive social valence to the pair-bonded partner. This may then permit the formation of reward/partner associated neurons in dCA1 through coactivation of CA2 and aminergic modulatory inputs to dCA1. Direct experimental investigation of this hypothesis, particularly using circuit-specific manipulations, will be important for determining how hippocampal-reward system connectivity contributes to pair-bond-related memories. Overall, our findings provide insight on the dual roles of CA1 and CA2 in social memory encoding and how their representations are modified as the pair-bond matures.

## Author Contributions

K. H., A.B., and S.A.S conceived the project and designed the experiments. K.H. collected and analyzed the data. O.G. scored tethered pair-bond videos. K.H., A.B., and S.A.S interpreted results and wrote the manuscript K.H., A.B., and S.A.S.

## Acknowledgments

This work was funded by National Institutes of Health grant MH104602 to S.A.S and a Ford Foundation Predoctoral Fellowship to K.H-P. We thank C. Gebhardt, M. Donegan, A. Oliva, and T. Tabachnik for technical support; S. Fusi, and. D. Aronov for critical discussions; R. Nguyen and L. Posani for helpful discussion on analysis. D. Franco for illustration of California mice.

## METHODS

### Subjects

All procedures were approved by the Columbia University Institutional Animal Care and use Committee and in accordance with NIH guidelines. *Peromyscus californicus insignis* (stock IS) were bred originally purchased from the *Peromyscus* Stock Center at the University of South Carolina and then a colony maintained at Columbia University. California mice were weaned at post-natal day 30. All mice were maintained under a 16:8 h light:dark day cycle and received food (PicoLab Rodent Diet 5053 for virgin animals, 5058 for breeders and male-female pairs) and water ad libitum. All mice were housed in 38.1 cm x 26.67 cm x 19.05 cm ventilated cages with corn bedding (NexGen Rat 900, Allentown). After weaning, all mice were housed in single-sex groups of 2–5 per cage. Mice used in experiments were between the ages of 3–11 months.

### Behavior

#### Tethered Pair-Bond Assay

The test was conducted in a new cage (38.1 cm x 26.67 cm x 19.05 cm) with corn bedding. Stimulus mice used were of opposite sex to the test mouse, one of which was the partner and the other a novel mouse. All pairs tested were co-housed for a week and video footage was recorded overnight to confirm mating.

Habituation was conducted in the first two days of the protocol. Test males and females were first habituated to a new empty cage for 20 min. Thereafter, test mice were habituated to the tether for another 20 min. Lastly, novel females and males were habituated to the tether for 20 min. On test day, the test mouse was habituated to a clean cage for 20 min. After this period, the test mouse was removed, and stimulus mice were placed in tethers in opposite corners of the cage for 20 min. After the stimulus mice were habituated, the test mouse was placed in the cage for 1 h. Time spent displaying affiliative contact and aggression frequency was quantified using manual scoring (Table 2). The test order and location of stimulus was counterbalanced between pairs.

**Table 1.**
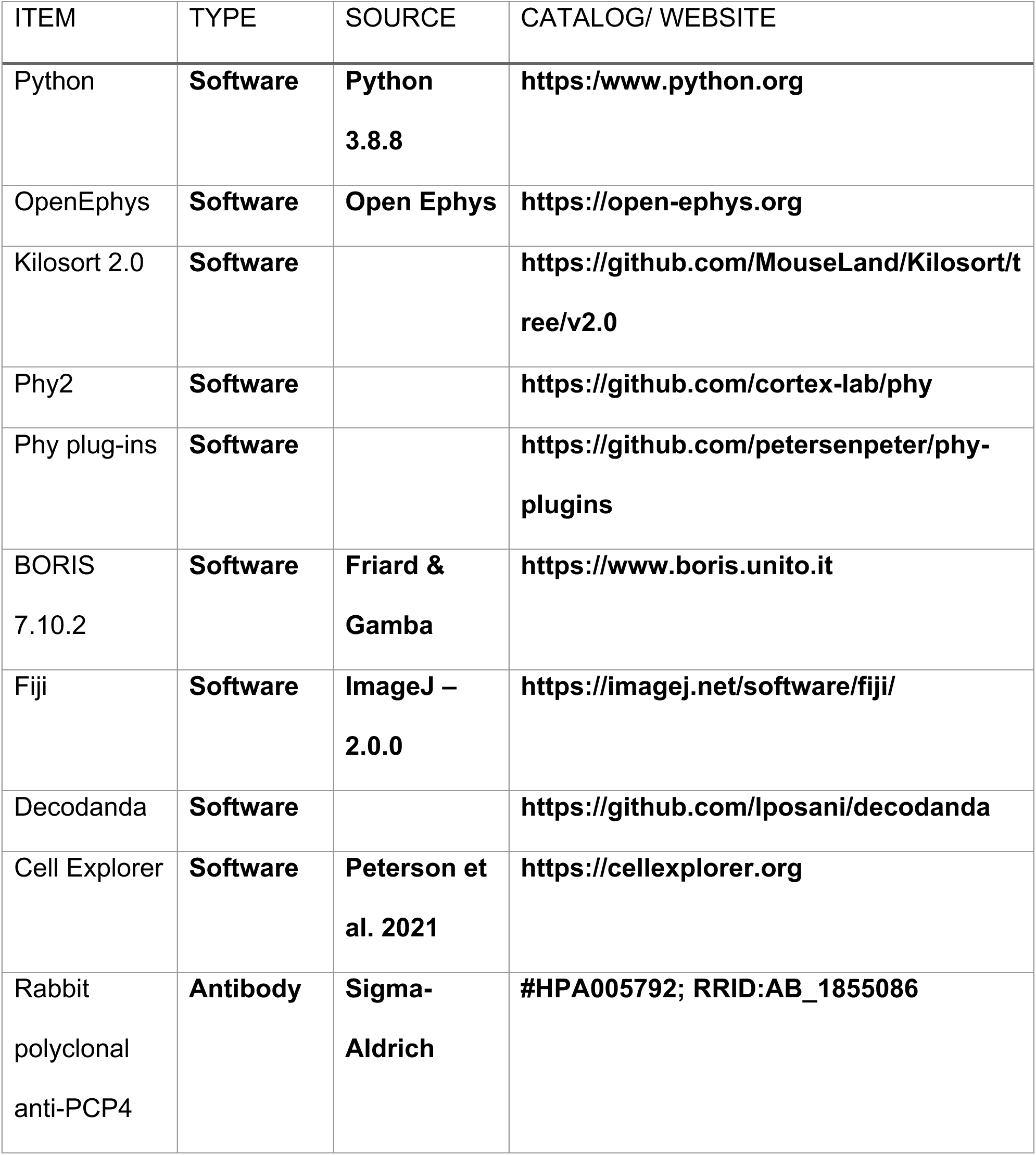
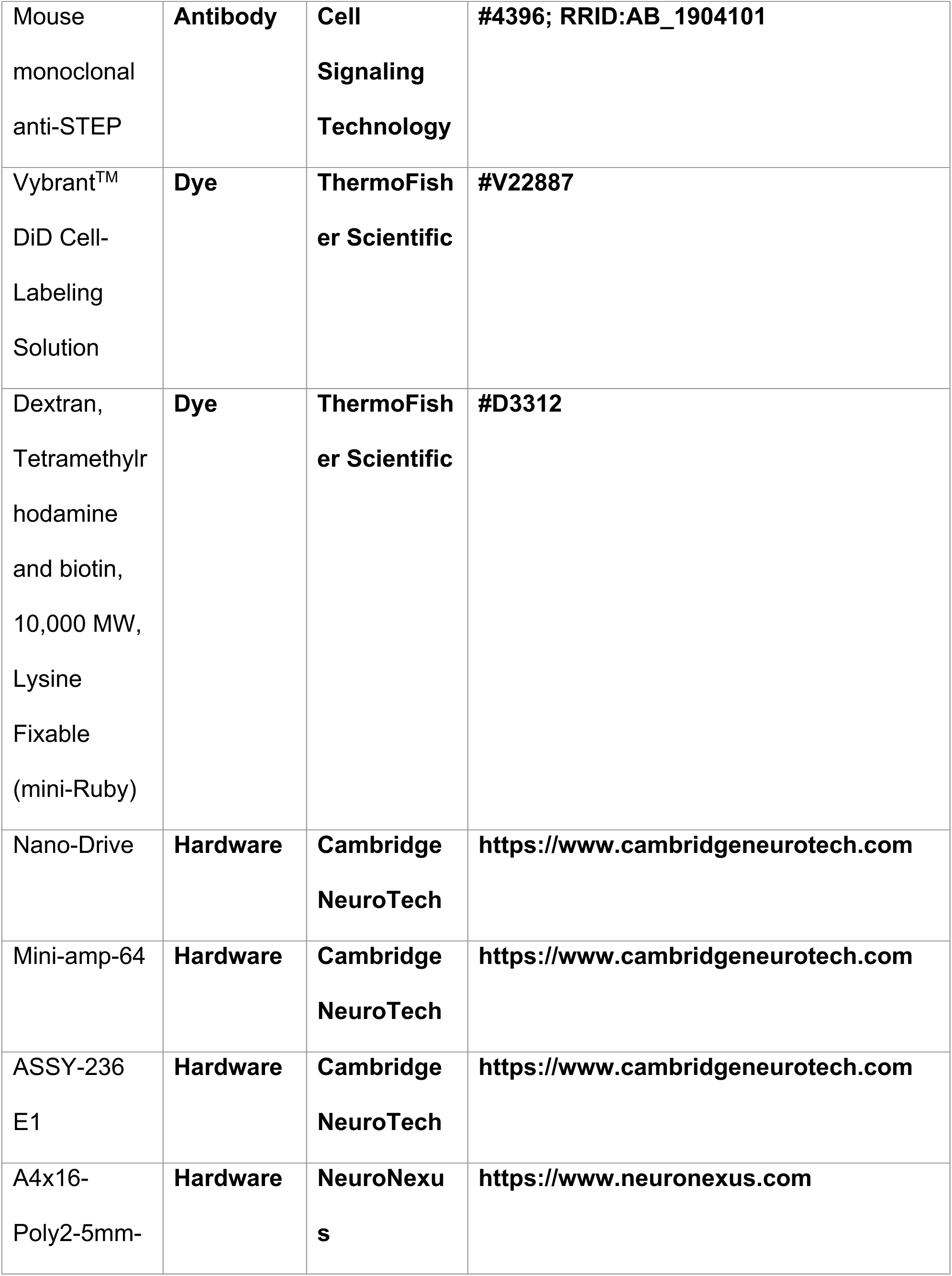
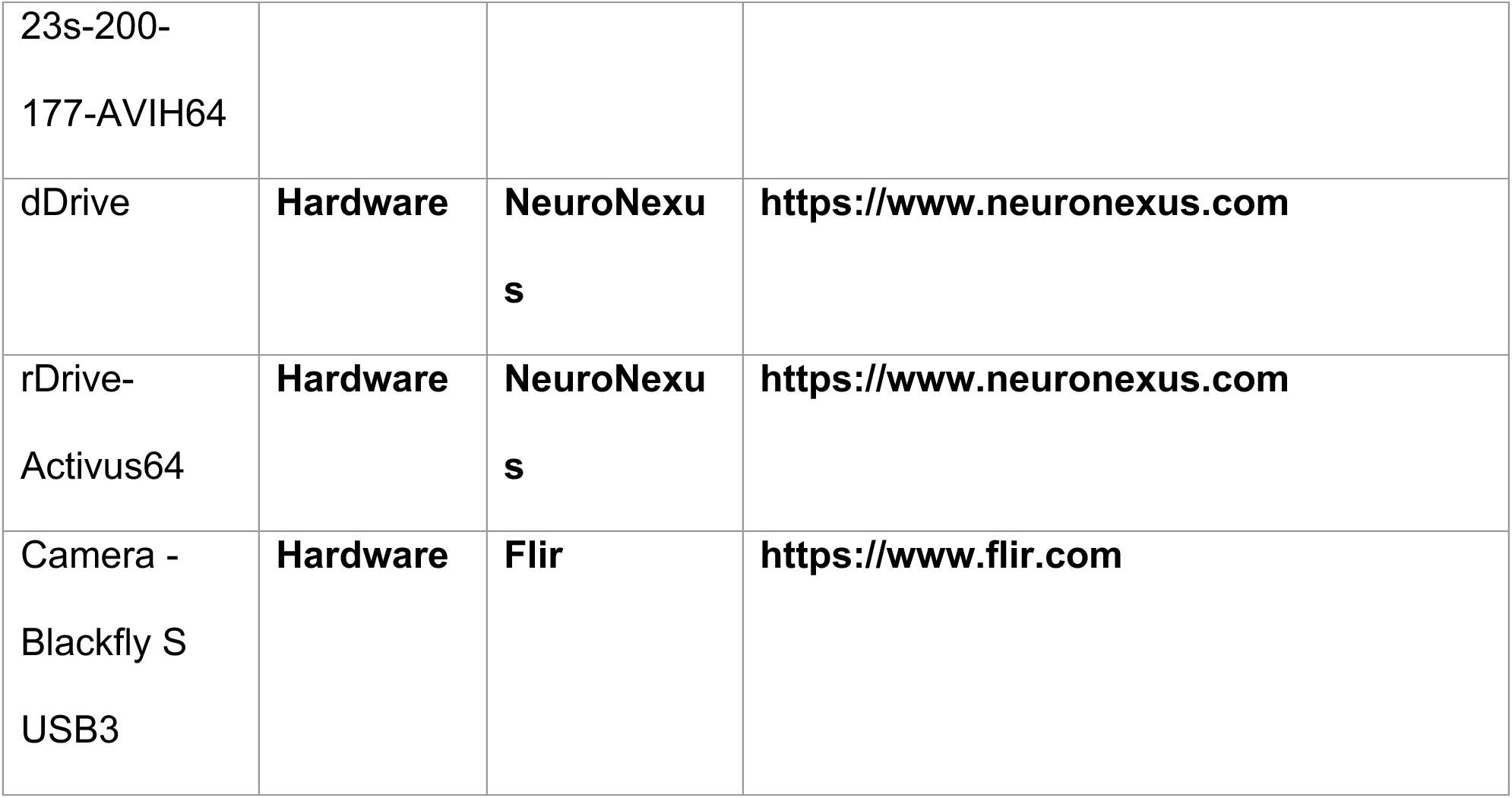

**Table 2.**

| **Behavior** | **Description** |
| --- | --- |
| **Affiliative contact** | A minimum of one-third of the animal's body length was required to be in direct physical contact |
| **Aggression** | A rapid acceleration of the test mouse toward the stimulus mouse, resulting in an aggressive behavior including, but not limited to, boxing, chasing, or biting. |

#### Pair-bond cups assay

Stimulus mice, one of which was the partner and the other novel, were of opposite sex to the test mouse. This assay consisted of five sessions. In session one, the test animal was placed in an empty 48.3 cm. x 48.3 cm x 38.1 cm box arena and allowed to explore for 20 min. In the next 10 min session, the test mouse explored the arena with two wire cups placed in the top-left and bottom-right corners of the arena (tall water bottles were placed on top of the wired cups to prevent climbing).

In session 3, stimulus mice were placed in the wire cups, and the test mouse was allowed to explore for 10 min. Placement of stimulus mice was counterbalanced between animals. After locations were cleaned with 70% ethanol and dried, the position of the stimulus mice was reversed for the 4^th^ session and test mouse was allowed to explore for another 10 min. Lastly, in the 5^th^ session, stimulus mice were removed, and the test mouse was alone in the area with empty cups for 20 min. The start of an investigation bout began when test mice were within a 5 cm threshold distance from nose to edge of cup and facing cup. Behavior was scored manually.

### CA2 Coordinate Identification

Mice were anesthetized using isoflurane and a small (<2 mm) hole was drilled in the skull with a sterile dentist’s drill above the target brain structure(s), which were located based on stereotaxic coordinates using bregma as a reference point. Needles pulled from borosilicate glass were used to deliver 10 nL of mRuby by injection through the drilled hole(s) in the cranium using the stereotactic manipulator and a nanopump. The mouse was then immediately perfused with 1 x PBS followed by 4% PFA. Brains were sliced, cut and imaged. Distance from dye to target location and bregma to lambda size for each mouse was collected. Using a dataset of coordinate estimations where mice had a bregma to lambda size ∼6.0 mm, an equation to estimate CA2 target anterior-posterior coordinate percentage was obtained using least squares regression: 12.3326 + 1.4025(Bregma-lambda size) = % → Multiply by Bregma-lambda size = – Anterio-Posterior coordinate. For example: 12.3326 + 1.4025(6.0) = 20.75 = –1.25 mm Anterio-Posterior coordinate. Other coordinates were consistent across mice, Medio-Lateral = 2.65 mm and Dorso-Ventral = –2.25 mm.

### Immunohistochemistry and Imaging

Mice were transcardially perfused using 1X PBS followed by 4% PFA in PBS. Brains were extracted and placed in 4% PFA overnight. Prior to sectioning, brains were washed for 1 h at room temperature (RT) in 0.3% glycine in PBS. Brains were sectioned at 40 µm using a Leica VT 1000S vibratome. Sections were then permeabilized and blocked for 4 h with a 5% goat serum, 0.5% Triton-X, and PBS solution at RT. Sections were incubated for two overnight periods with primary antibodies in a 5% goat serum, 0.1% Triton-X solution, and PBS solution. Sections were then washed with PBS 3 times for 10 min at RT. Goat secondary antibodies were then applied in a 5% goat serum, 0.1% Triton-X solution, and PBS solution for 4 h at RT. Lastly, DAPI (ThermoFisher Scientific, #D1306) was applied for 20 min at 1:1000 dilution in PBS at RT. Sections were then mounted and covered using Fluoromount (Sigma-Aldritch). Primary rabbit anti-PCP4 (1:300, #HPA005792, Sigma-Aldrich) and mouse IgG1 anti-STEP (1:1000, #4396, Cell Signaling Technology) followed by secondary goat anti-rabbit (1:500, #A-11008, Invitrogen) or goat anti-mouse IgG1 (1:500, #A-21121, Invitrogen) conjugated to Alexa 488 was used to identify CA2.

### Extracellular Probe Implantations

To ensure exploratory test mice for probe implantation, mice were prescreened in a 5 min open field test to assess exploratory behavior. Mice that demonstrated constant exploration of the entire open field arena throughout the 5 min test were chosen for implantation. Mice were administered carprofen (5 mg/kg) 1 h prior to surgery. Mice were anesthetized using 0–5% isoflurane. Bone was scratched slightly using a sharp forceps to allow for better adhesion of the dental cement. Metabond Enamel Etchant Gel was applied for 20 seconds, rinsed and dried, followed by Metabond Universal Dentin Activator for 30 seconds, rinsed and dried. Two small stainless screws were driven into the skull to anchor the implant and were covered with dental cement. A small (∼0.5 mm) craniotomy was made above the cerebellum, in which a 99.9% silver ground wire was placed. Dental cement was placed around the skull attached with copper mesh around the head of the animal. A copper mesh hat served to protect the implant and decrease electrical noise. Silicone probes used had a four-shank design (Cambridge NeuroTech: ASSY-236 E1, NeuroNexus: A4×16-Poly2-5mm-23s-200-177-AVIH64). Probes were adhered to a microdrive (Cambridge NeuroTech: Nano-Drive, NeuroNexus: dDrive, rDrive-Activus64) to allow for vertical manipulation of probes until target location was reached. A few drops of DiD Cell-Labeling Solution (#D3312, ThermoFisher Scientific) was applied above channels on the probe to identify probe location at the end of experimentation. Another craniotomy (1 mm x 0.5 mm) was drilled above the intended cortical or subcortical targeted regions. The electrode was driven into the brain slowly till reaching outside the hippocampus (DV = 1.80 mm) and then the microdrive was secured with dental cement. More cement was used to build a stronger, stable implant around and above the head of the animal. Buprenorphine XR (3.25 mg/kg) was administered 12 hours post-surgery.

### Recordings

Mice were allowed to recover for four days before being handled. After the four-day period, probes were lowered slowly 50–150 µm a day until the hippocampal pyramidal layer was reached, as determined by single-unit spike activity and local field potential (LFP) patterns (theta oscillation and sharp-wave ripples). Electrophysiological recordings were amplified and digitized using the OpenEphys GUI. Data was collected at a rate of 30 kHz. OpenEphys files were converted to binary files and a thresholded for 6 standard deviations prior to being semi-automatically spike sorted using Kilosort 2.0 (https://github.com/MouseLand/Kilosort/tree/v2.0). Units were visually inspected using Phy2 and were accepted, split, merged, or eliminated. Custom designed plug-ins were also used to characterize well isolate units (https://github.com/petersenpeter/phy-plugins). Each unit’s waveform shape, stability, and autocorrelograms were assessed to classify cells as either inhibitory or excitatory and confirmed using Cell Explorer. Implanted mice were perfused, and brains were collected, sliced, and imaged at the conclusion of recordings to identify the recording site.

### Single-cell responses

#### Activation and inhibition selectivity

Neurons were identified as activated or inhibited using a permutation-based analysis comparing firing rates during investigation (post-onset) of stimuli versus baseline (pre-onset) for each stimulus separately. For each neuron, firing rate data across sessions was combined and the observed mean difference between investigation bouts and baseline was computed. Bouts were binned by 250 ms. The number of bouts per condition (pre- and post-) was balanced to ensure equal representation for each neuron. The amount of bout bins used in the permutation was determined by the minimum number of bins in session/pre- or post-onset/stimulus condition to ensure equal representation of each session/condition. If a session/condition only contained <3 bouts, this data was duplicated to allow for 1000+ unique shuffles (occurring 7/70 experiments).

To generate a null distribution, bout-level stimulus labels were randomly shuffled 1000 times while preserving the bout and session structure. For each shuffle, the mean firing rate difference between conditions was recalculated, creating a distribution of permuted differences under the null hypothesis of no stimulus preference. A neuron was classified as inhibited if the observed difference fell in the lower 2.5% of the null distribution (i.e. greater firing during partner investigation). Activated cells fell in the upper 2.5% of the distribution. Neurons whose observed difference did not meet either threshold were labeled as not responsive.

#### Stimulus selectivity

Neuron selectivity for stimuli was determined using a permutation-based analysis comparing firing rates during investigation of partner versus novel conspecifics. For each neuron, firing rate data across sessions was combined and the observed mean difference between partner and novel investigation bouts was computed. We started collecting bins 1 s before start of stimulus investigation periods. Bouts were binned by 250 ms. The amount of bout bins used in the permutation was determined by the minimum session/stimulus condition. For example: Session 1 with partner contained 50 sec of data, while session 1 with novel had 120 sec, session 2 had 102 sec with partner and session 1 contained 95 sec. Session 1 data length was used to determine the amount of data obtained from the other conditions. The size of the bouts by number of bins set by minimum session/stimulus condition was matched to the other conditions. If a session/condition only contained <3 bouts, this data was duplicated to allow for 1000+ unique shuffles (occurring 7/70 experiments). The number of bouts per stimulus type was balanced to ensure equal representation for each neuron.

To generate a null distribution, bout-level stimulus labels were randomly shuffled 1000 times while preserving the bout and session structure. For each shuffle, the mean firing rate difference between conditions was recalculated, creating a distribution of permuted differences under the null hypothesis of no stimulus preference. A neuron was classified as novel-selective if the observed difference fell in the lower 2.5% of the null distribution (i.e. greater firing during partner investigation). Partner-selective cells fell in the upper 2.5% of the distribution. Neurons whose observed difference did not meet either threshold were labeled as having no preference.

### Data labeling

Time periods of active exploration were identified when animals were tested in the pair-bond cups (PBC) task. These periods were defined when the mouse was within 5 cm distance to the edge of cups and facing an angle of 0-90 degrees towards the center of the cup (0 being when the mouse was directly facing the center of the cup). These periods were then broken up into 250 ms firing rate bins. Each of these bins were labeled as interactions with either partner or novel mouse and location of the cup, left or right. The PBC task contained two sessions resulting in four conditions: [Partner, Left], [Novel, Left], [Partner, Right], and [Novel, Right].

### Decoding

A support Vector Machine with a linear kernel was implemented using the decoding package Decodanda (https://github.com/lposani/decodanda). Decoding analysis used pseudo-population activity from all animals tested, with 75% used to train the classifier and 25% used to test the decoding performance. A 10-fold cross validation was implemented throughout the analysis. Null models were generated by shuffling the decoding variable (mouse or location) and neural data. The decoding procedure (resampling + cross validation) was repeated for 25 null iterations to test statistical significance. Final decoding performance was computed as the average classification accuracy across all cross-validation folds. Null distributions were used to compute statistical significance of observed performance. A p-value was obtained from the z-score of the performance computed on data compared to the distribution of n null-model values: z = (μ -< μ>)/ σ. In order to compare decoding performance across days or regions, we ensured the same number of cells and same number of time bins were used with each decoding procedure. Animals that explored both stimulus mice in both session for a minimum of 4 s were used in analysis.

### Statistics

Statistical significance was defined as *, p<0.05; **, p<.01; ***, p<0.001.

#### Tethered Pair-Bond, Pair-Bond Cups

To test significance difference between groups (i.e. partner vs. novel), we ran paired t-test for total time spent investigating stimuli. In samples <10, we ran a Wilcoxon signed-rank test. Discrimination index (DI) was calculated based on novel and partner exploration times:

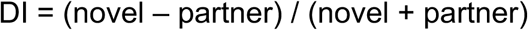

To test significance, index scores were tested in one-sample t-test compared to 0.

#### Average Firing rates & Stimulus average firing rate response

To test for differences in firing rates, we implemented an independent samples t-test for cross region comparisons. For comparison of firing rates during stimulus investigations across regions, we used a linear mixed-effects model with region as a between-subject factor, stimulus as a within-subject factor, and subject identity (ID) included as a random effect. Fixed effects were tested using Wald z-statistics. Significant interactions were followed up with post hoc comparisons, including paired t-tests for within-region comparisons and Welch’s t-tests for between-region comparisons. A Bonferroni correction was applied to control for multiple comparisons.

#### Cell selectivity analysis

To test for significant differences in distributions of cell selectivity categorization for each region, we implemented a two-sided Fisher’s Exact Test with post hoc pairwise comparisons. A Bonferroni correction was used to control for multiple comparisons across categories.

To compare stimulus selectivity distribution between regions on day 0, we ran a Fisher’s Exact test. To identify which specific category contributed to regional difference, we performed post hoc pairwise comparisons using 2×2 Fisher’s exact tests (each category vs. all other categories). A Bonferroni correction was used to control for multiple comparisons across categories.

#### Stimulus-selective responses

To test for significance of differences between partner-(or future partner) and novel-selective cell responses, we ran a paired t-test comparing mean firing rates and changes from baseline.

#### Region and day decoding comparisons

To assess how decoding performance varied as a function of region and subsample value, we fit a linear regression model with decoding performance as the dependent variable and region (two levels), subsample value (continuous), and their interaction as predictors:

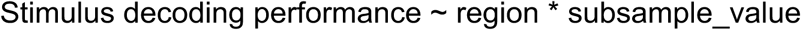

The region × subsample value interaction tested for differences in slope between regions. Model effects were evaluated using Type II analysis of variance (ANOVA) F-tests.

## SUPPLEMENTAL FIGURES

**Supplemental figure 1.**
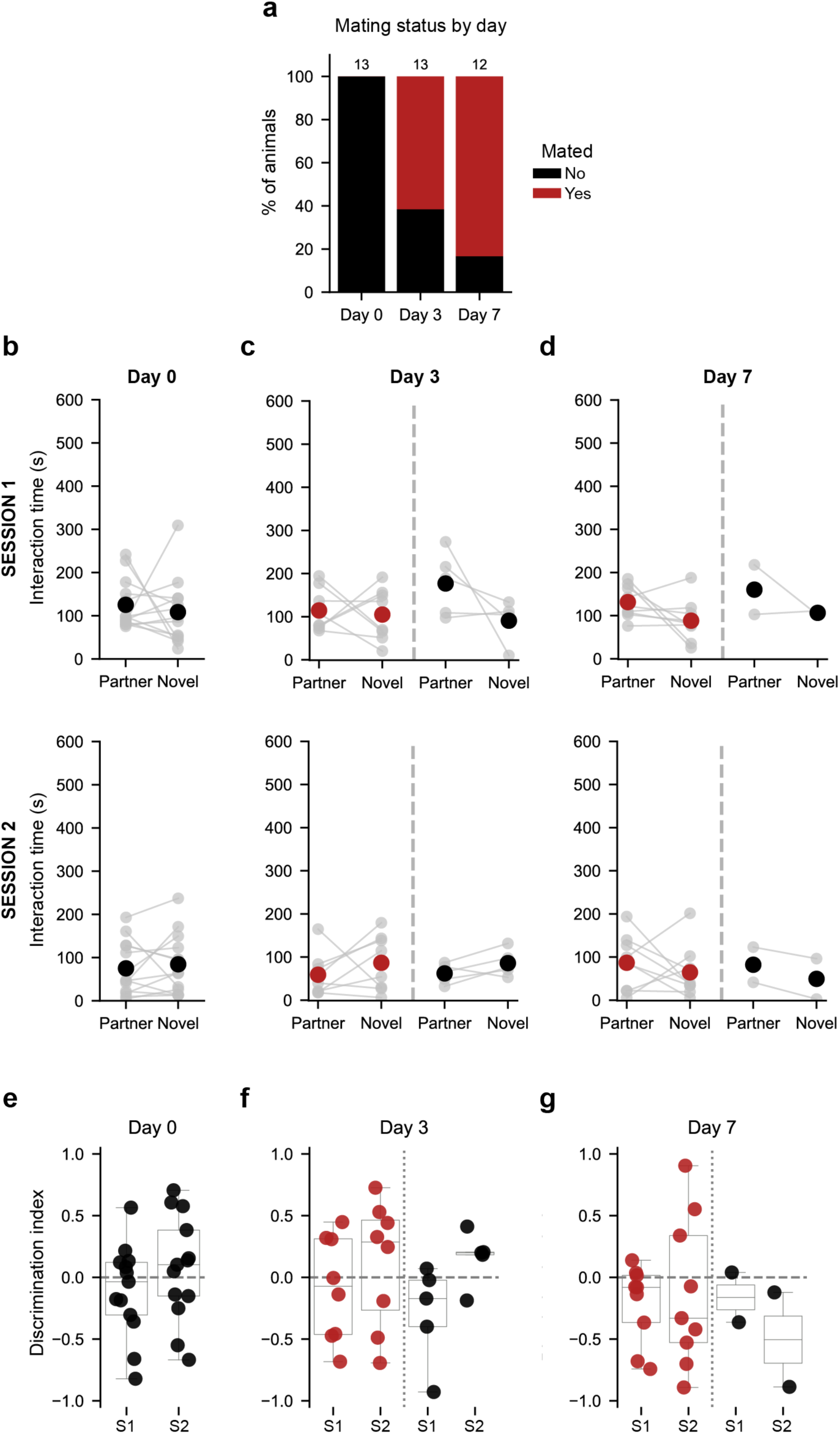
Animal status and interaction times during pair-bond cups task. **a)** Mating status across recording days: mated (red) and not mated (black). By day 3, 8 out 13 mice mated. On day 7, 10 out of 12 mice mated. **b)** Session 1 (top row), session 2 (bottom row). Day 0, Test mice spent an equal amount of time exploring both stimulus mice. Session 1 and 2: Paired t-test = not significant (ns), n = 13. **c)** Day 3, No significant difference in interaction times observed (paired t-tests; Mated n = 8, Not Mated = 5). **d)** No difference in interaction time on day 7 (paired t-tests; Mated n = 9, Not Mated n = 2). **e - g)** Discrimination index: (novel-partner)/(novel+partner) interactions times during sessions. No exploration difference was found across days. Boxplots indicate median and interquartile range.

**Supplemental figure 2.**
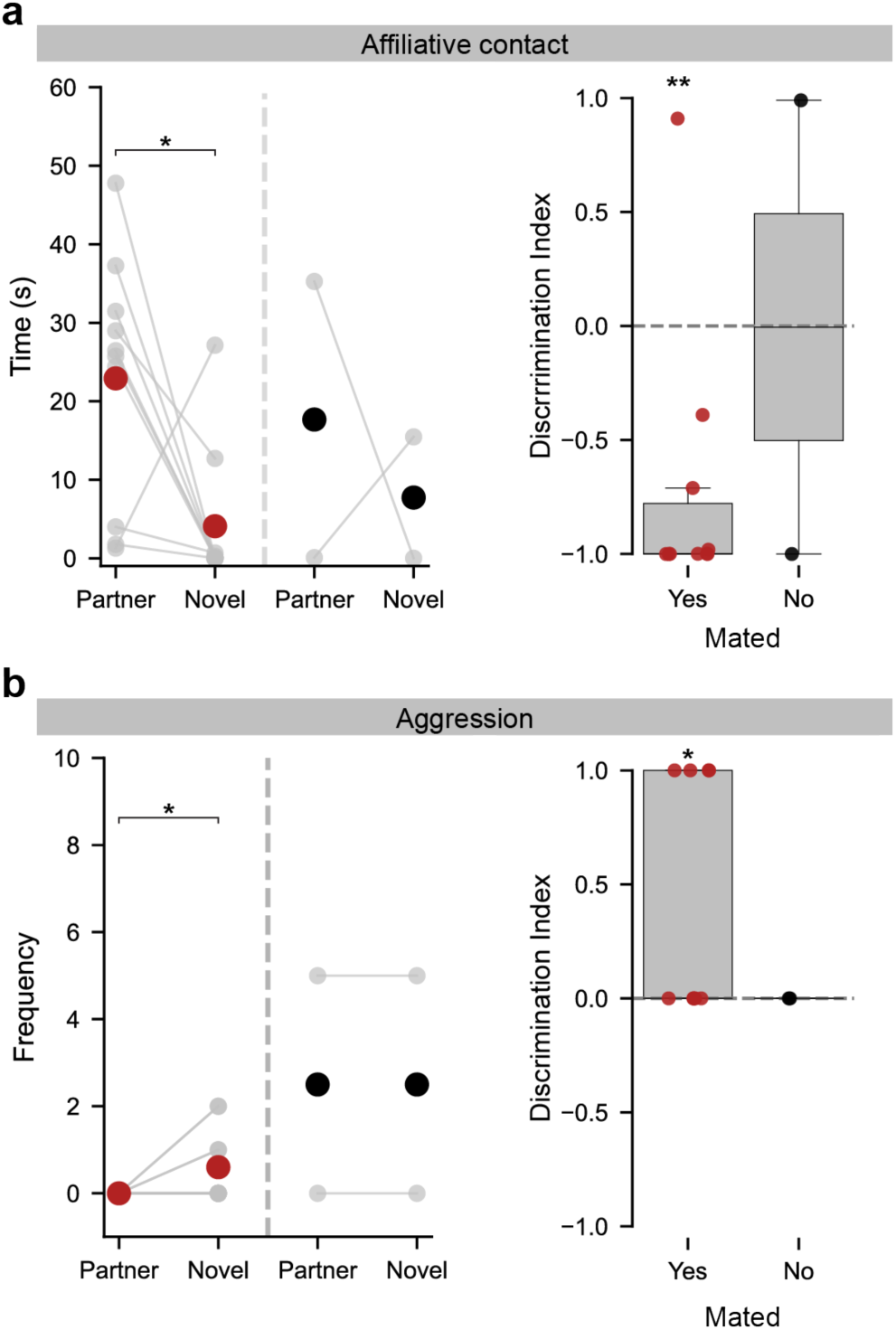
Implanted mice exhibit selective affiliative contact and aggression to co-housed partner. Mated (red), Not mated (black). **a)** (Left). Males displayed greater affiliative contact towards partner compared to novel female. Mated: Paired t-test, t(9) = 2.823, *p* = 0.0199, n=10. (Right) Affiliation discrimination index for partner versus novel mouse interaction times. Discrimination Index = (novel - partner)/(novel + partner) interaction times during session. Mated: One-sample t-test against 0, t(9) = −3.743, *p* = 0.0046 (n=10). **b)** (Left) Mated animals displayed a trend to increased aggression towards novel stimulus compared to partner. Mated: Paired t-test, t(9) = −2.250, *p* = 0.0510, n = 10. (Right) Aggression discrimination index for novel versus partner. Discrimination index = (novel – partner)/(novel + partner) number of attacks per session. Mated: One-sample t-test against 0, t(9) = 2.449, p = 0.0368, n=10. Boxplots indicate median and interquartile range. \**p* < 0.05, \*\**p* < 0.01.

**Supplemental figure 3.**
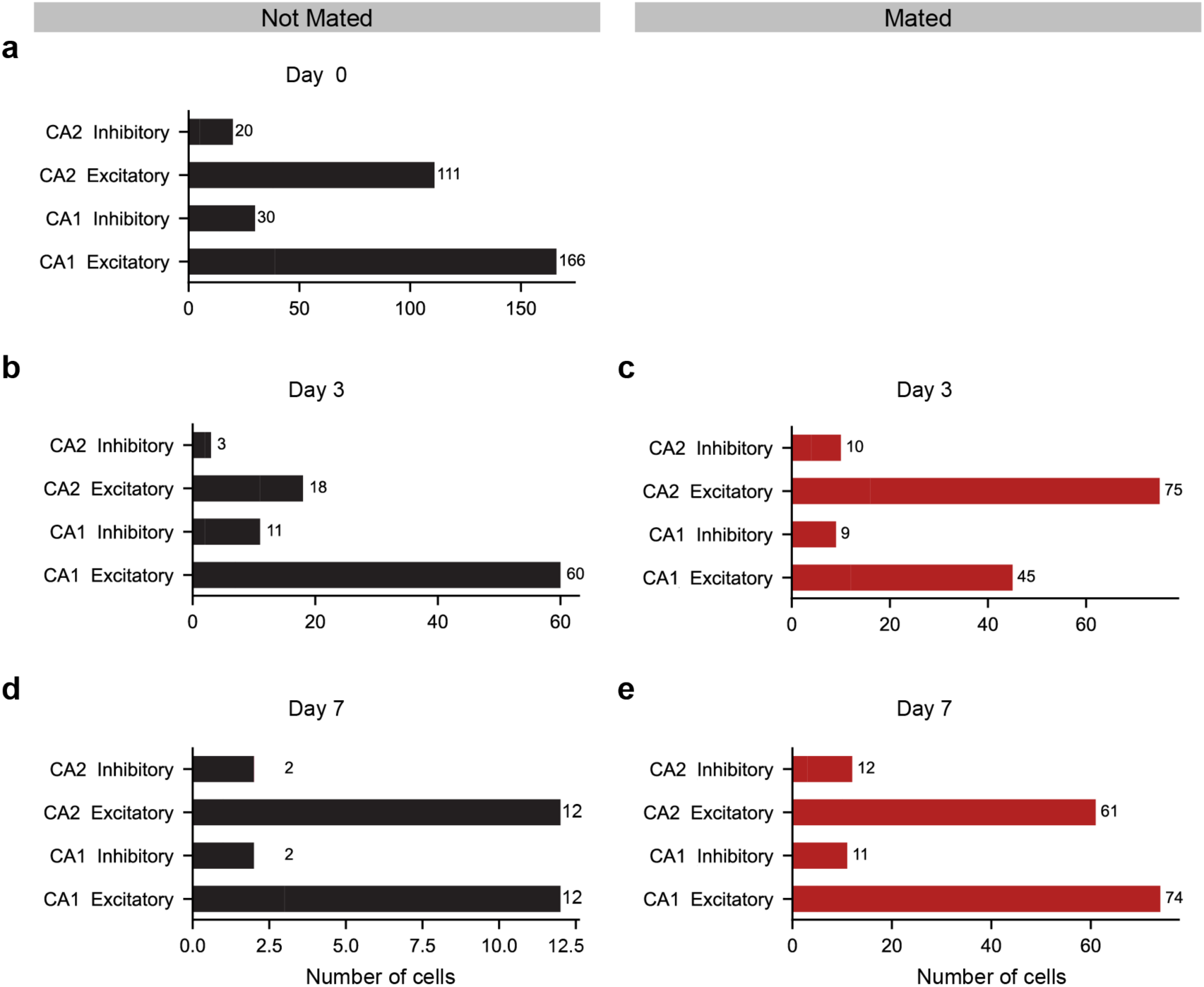
Excitatory and inhibitory cell counts across days and mating status. Recordings were obtained from both CA1 and CA2 regions of the hippocampus to enable regional comparisons during pair-bond formation. Bars show total numbers of excitatory and inhibitory neurons recorded from all animals in a given region on a given day. Not mated (black), Mated (red). **a)** Day 0, Not Mated: n = 9 mice for CA1 and 8 mice for CA2. **b)** Day 3 – Not Mated: n = 5 mice for CA1, 3 mice for CA2. **c)** Day 3 – Mated: n = 4 mice for CA1, 5 mice for CA2. **d)** Day 7 – Not Mated: n = 2 mice for CA1 and 2 CA2. **e)** Day 7 - Mated: n = 5 mice for CA1, 4 mice for CA2.

**Supplemental figure 4.**
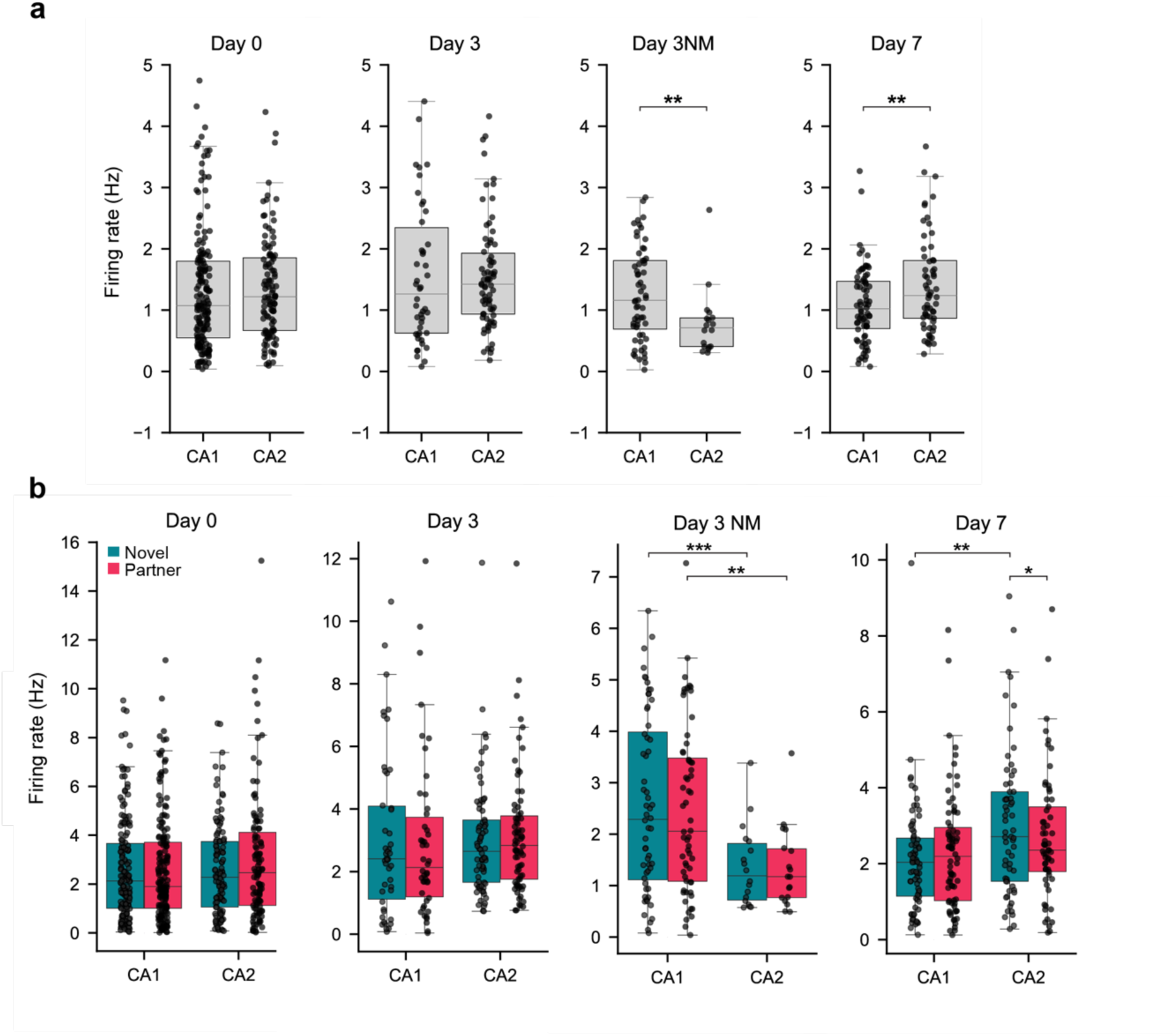
Dependence of average CA1 and CA2 excitatory cell firing rates across days and during interactions with stimulus mice. **a)** Average firing rates (FR) during the two sessions across regions. On day 3 for the NM group, CA1 exhibited a significantly greater average FR than CA2 (Independent samples t-test, t(76) = 2.911, *p* = 0.0060). On day 7, average FR was significantly greater in CA2 (Independent samples t-test, t(133) = −2.635, *p* =0.0096). **b)** Average FR during interactions with indicated stimulus mice. Data averaged from both sessions for each region. No significant difference in firing rate was observed between CA1 and CA2 or during interactions with partner compared to novel stimulus mouse except as follows. On day 3 in the NM group, CA1 firing rate was significantly greater than CA2 firing rate during interactions with either novel or partner mouse. A linear mixed-effects model revealed a significant effect of region (z= −2.952, *p*= 0.003), followed by Welch’s t-test with Bonferroni correction applied for multiple comparisons (Novel: t(64.05) = 4.196, *p* = 0.0003, Partner: t(64.41) = 4.017, *p* = 0.007). On day 7, CA2 FR was greater during interactions with the novel mouse compared to the partner. Linear mixed-effects model revealed a significant effect of region (z = 3.341, *p*=0.001) and region x stimulus interaction (z = −2.962, *p* = 0.003), followed by paired t-test with Bonferroni correction applied for multiple comparisons (t (60) =2.805, *p*=0.006). Firing rate during interactions with the novel mouse was also greater in CA2 than CA1(Welch’s t-test, t (108.03) =-3.191, *p* = 0.001 Bonferroni adjusted). Boxplots indicate median and interquartile range. \**p* < 0.05, \*\**p* < 0.01, \*\*\**p* < 0.001.

**Supplemental figure 5.**
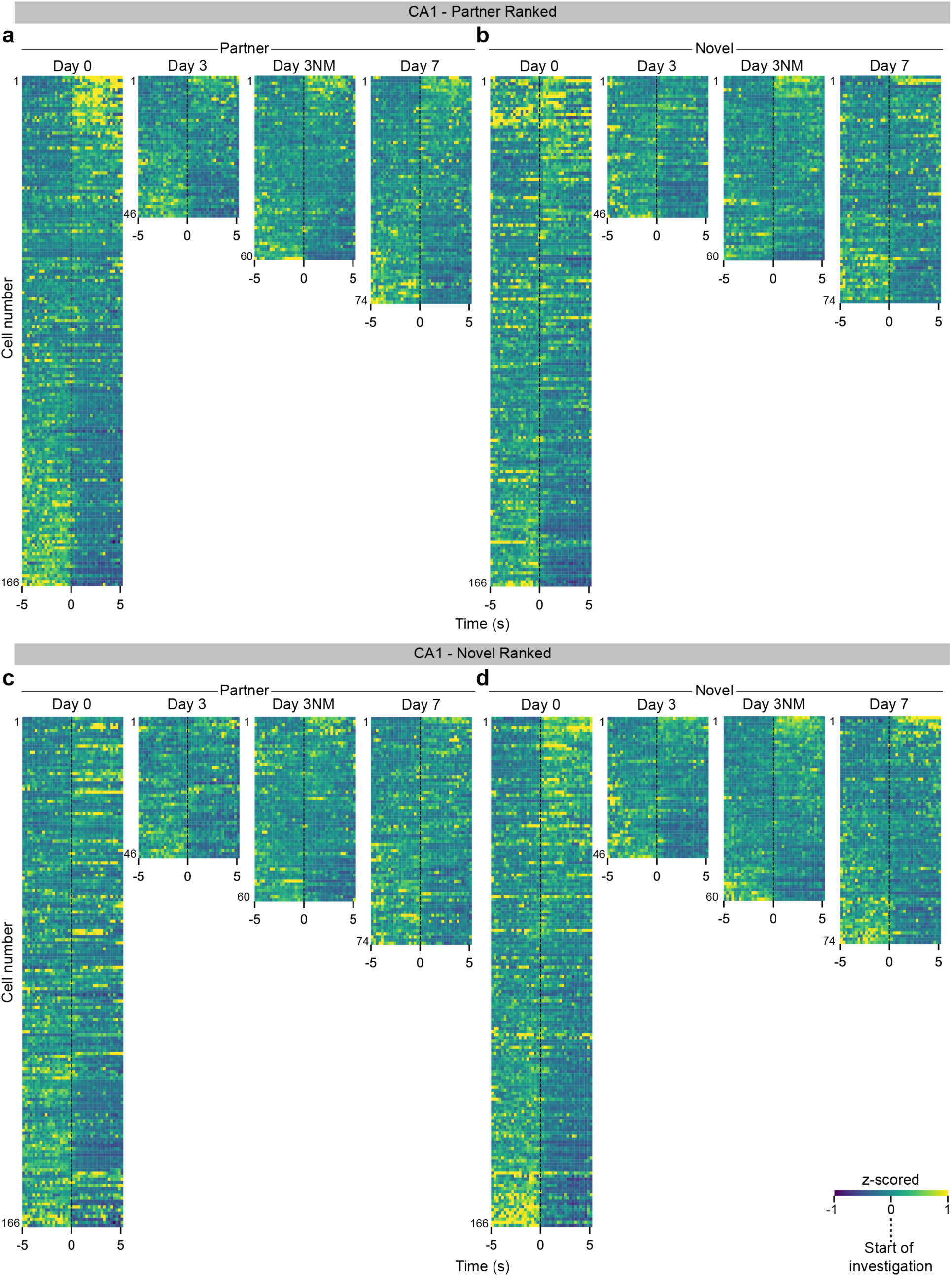
CA1 excitatory cell firing rates during interactions with partner and novel mouse on different days. Z-scored excitatory firing rates aligned to the onset of stimulus investigation (time 0). Each row shows the color-coded Z-scored activity of a neuron from the indicated region, starting 5 seconds before onset of investigation to 5 seconds after. Ranking was determined by the magnitude of the change in firing rate (FR) during the interaction relative to baseline, calculated as (FR_after – FR_before), averaging over windows t= 5 to 1 s before interaction and t= 0 to 5 s after start of interaction. **a)** CA1 excitatory activity to partner stimuli, ranked by partner responses. **b)** CA1 excitatory activity to novel stimuli, ranked by partner responses. **c)** CA1 excitatory activity to partner stimuli, ranked by novel responses. **d)** CA1 excitatory activity to novel stimuli, ranked by novel responses. CA1 cells, n = 166, 46, 60, 74 on day 0, 3, 3 NM, and 7 respectively. NM = Not Mated. Partner-ranked data from day 0 and 7 are same as depicted in Figure 3.

**Supplemental figure 6.**
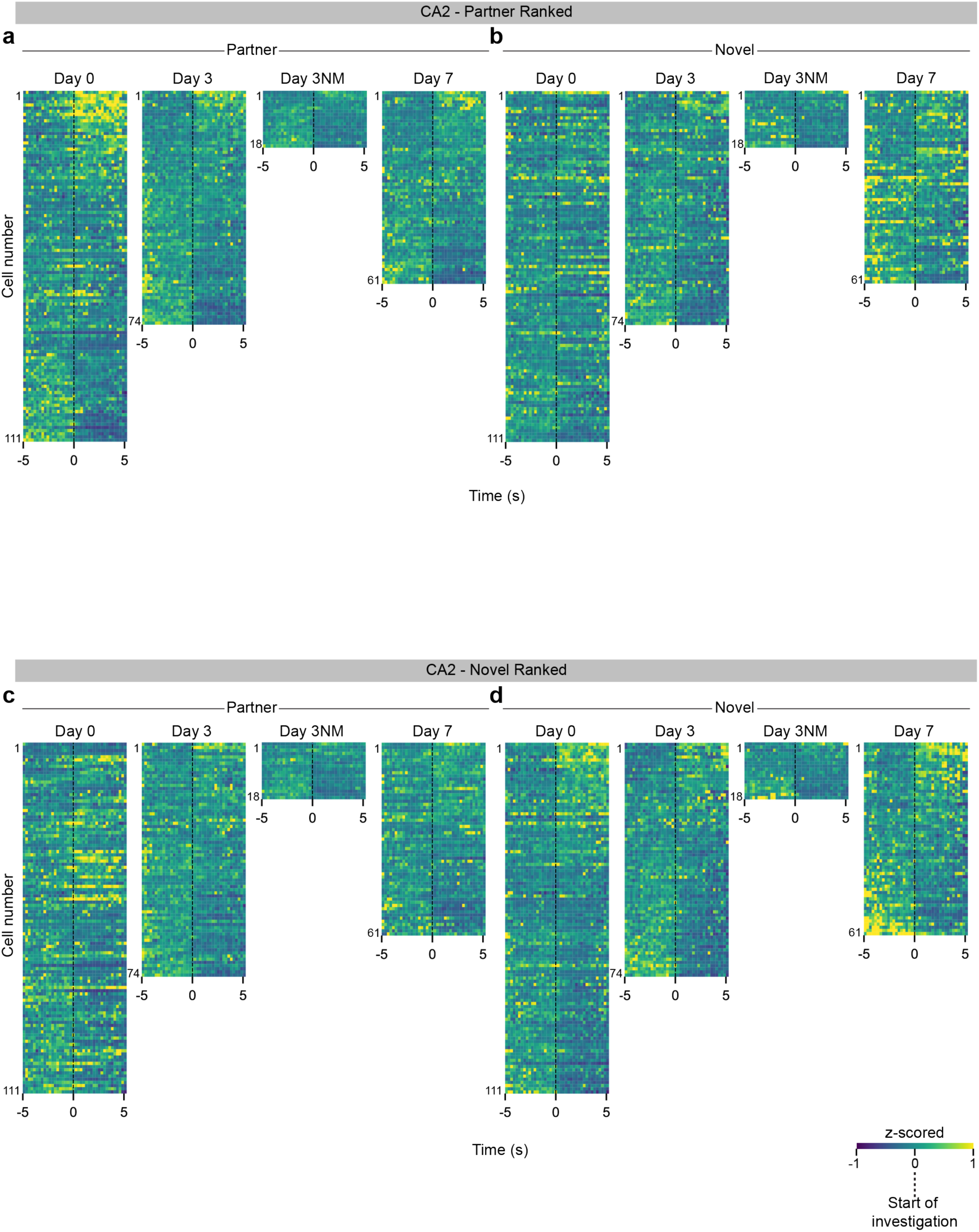
CA2 excitatory cell firing rates during interactions with partner and novel mouse on different days. Data displayed as in Supplemental Figure 5. **a)** CA2 excitatory activity to partner stimuli, ranked by partner responses. **b)** CA2 excitatory activity to novel stimuli, ranked by partner responses. **c)** CA2 excitatory activity to partner stimuli, ranked by novel responses. **d)** CA2 excitatory activity to novel stimuli, ranked by novel responses. CA2 cells, n = 111, 75, 18, 61 on day 0, 3, 3 NM, and 7, respectively. NM = Not Mated. Partner-ranked data from day 0 and 7 are same as depicted in Figure 3.

**Supplemental figure 7.**
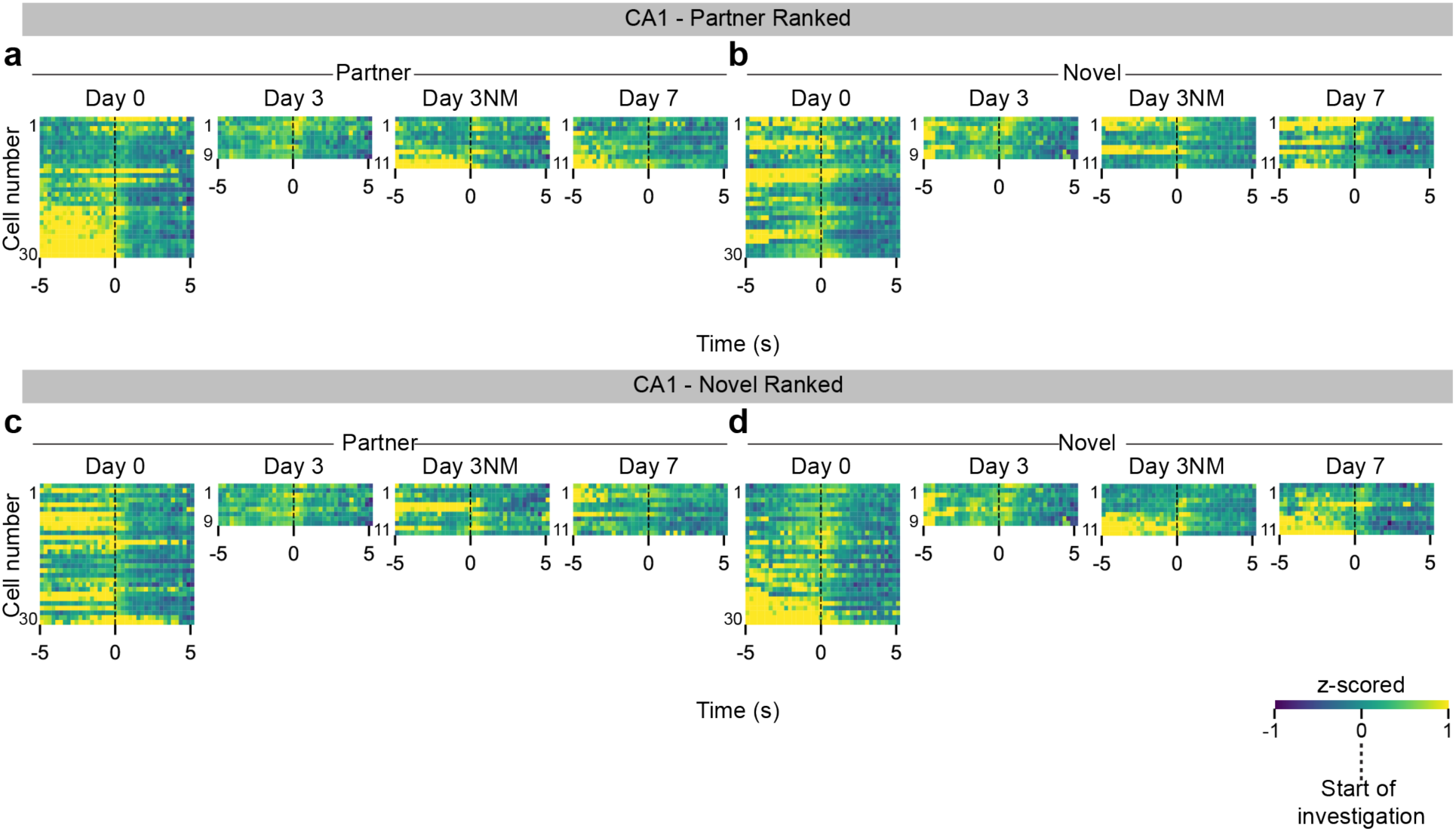
CA1 inhibitory cell firing rates during interactions with partner and novel mouse on different days. Data displayed as in Supplemental Figure 5. **a)** CA1 inhibitory cell activity to partner stimuli, ranked by partner responses. **b)** CA1 inhibitory cell activity to novel stimuli, ranked by partner responses. **c)** CA1 inhibitory cell activity to partner stimuli, ranked by novel responses. **d)** CA1 inhibitory cell activity to novel stimuli, ranked by novel responses. CA1 cells, n = 30, 9, 11, 11 on day 0, 3, 3 NM, and 7 respectively. NM = Not Mated.

**Supplemental figure 8.**
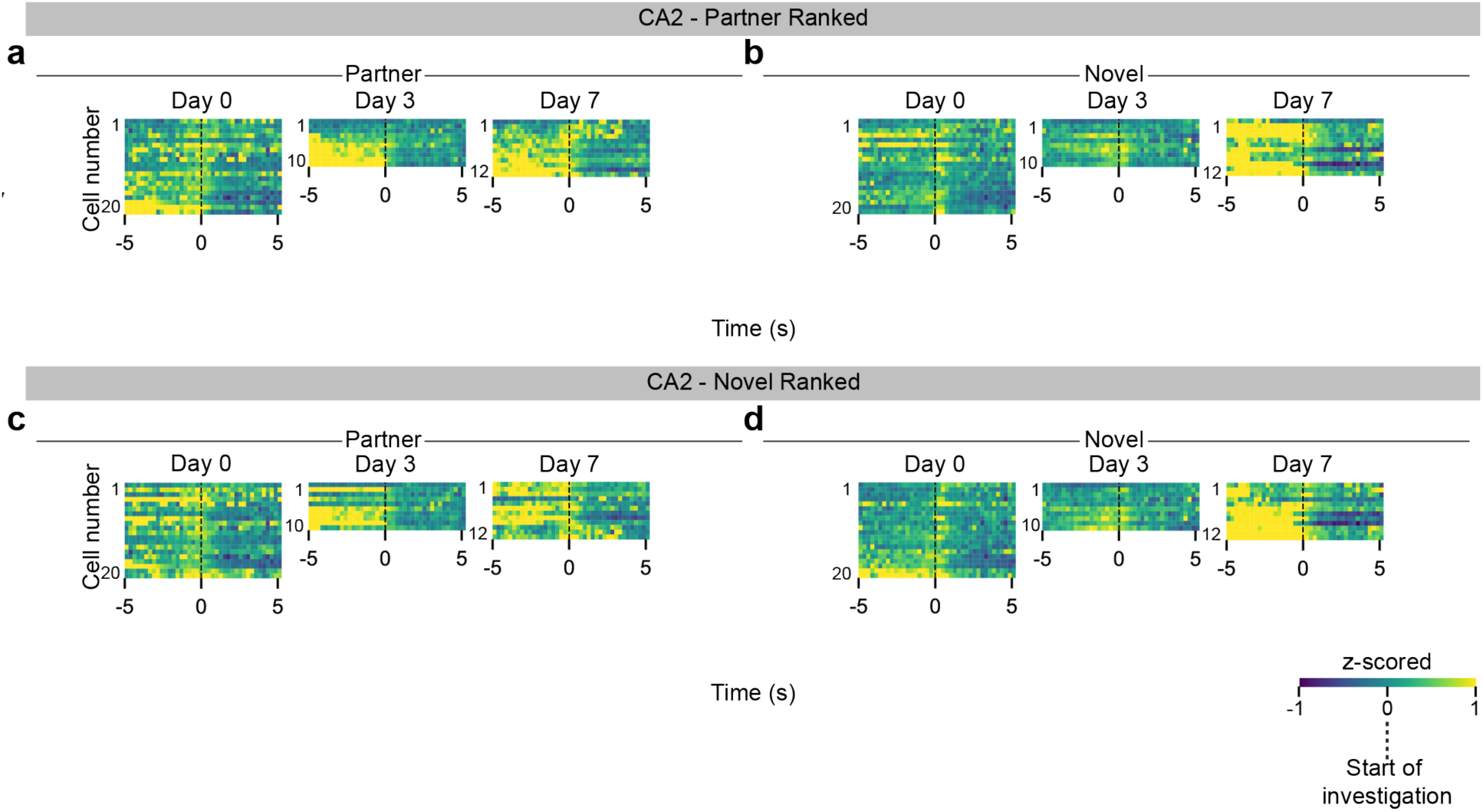
CA2 inhibitory cell firing rates during interactions with partner and novel mouse on different days. Data displayed as in Supplemental Figure 5. **a)** CA2 inhibitory cell activity to partner stimuli, ranked by partner responses. **b)** CA2 inhibitory cell activity to novel stimuli, ranked by partner responses. **c)** CA2 inhibitory cell activity to partner stimuli, ranked by novel responses. **d)** CA2 inhibitory cell activity to novel stimuli, ranked by novel responses. CA2 cells, n = 20, 10, 12 on day 0, 3, and 7 respectively. NM = Not Mated.

**Supplemental figure 9.**
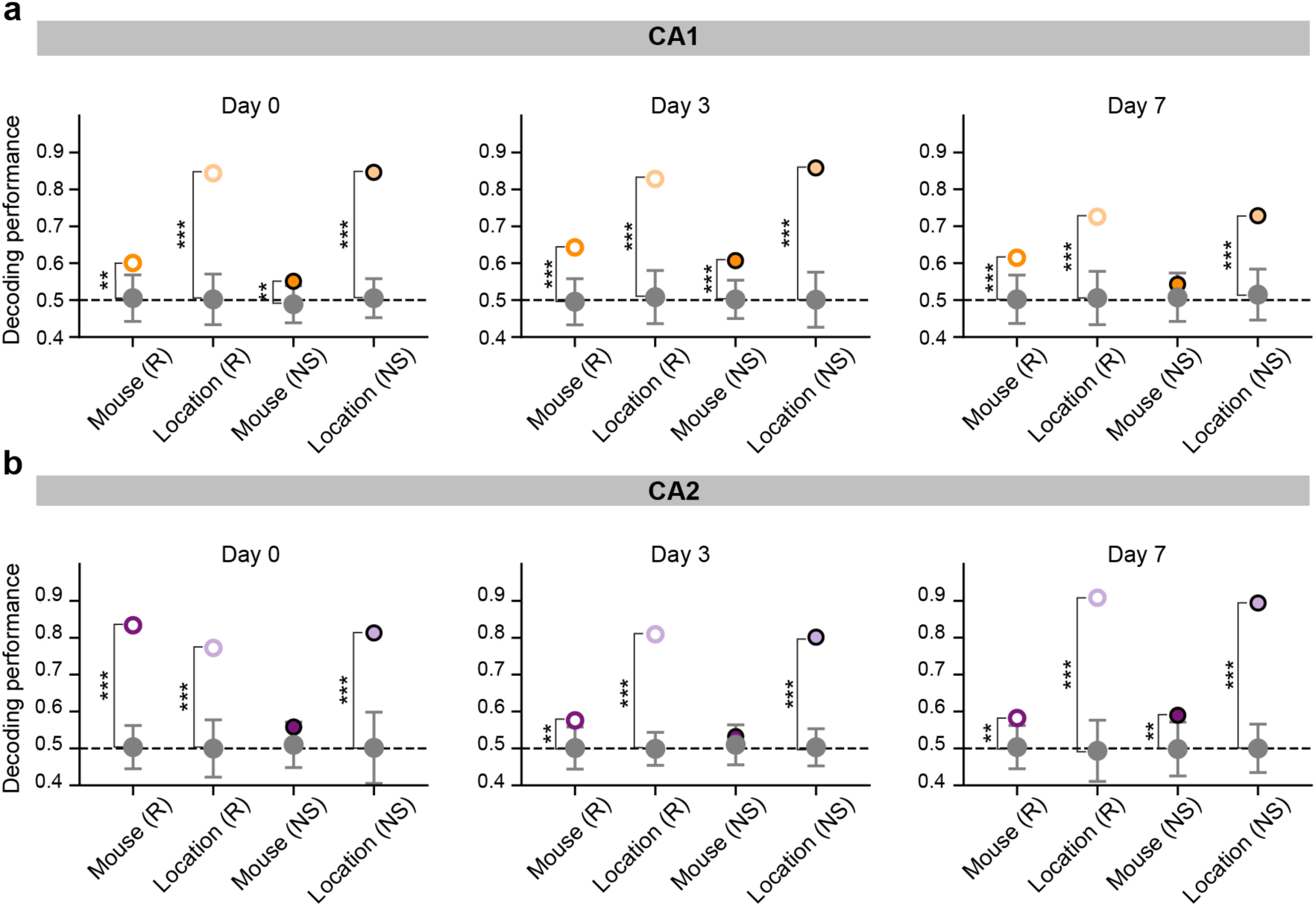
Effect of stimulus-preferring-cell activity on mouse and location decoding accuracy. Decoding performance compared when either mouse-preferring cells are removed, labeled with “NS” (filled circles), or an equal number of randomly selected cells are removed, “R” (empty circles), were removed from the pseudo-population. Darker circles represent mouse, while lighter circles represent location. Null model chance decoding in grey. **a)** CA1 results. Mouse and location decoding performance remained above chance when either stimulus-preferring-cells or random cells were removed, except on day 7 when decoding of mouse identity was reduced to chance levels absent the mouse-preferring cells. Day 0: mouse(R) decoding= 0.60, null model = 0.51±0.03; location(R) decoding = 0.84, null model = 0.50±0.03; mouse(NS) decoding= 0.55, null model = 0.49±0.03; location(NS) decoding = 0.85, null model = 0.51±0.03. (Pseudo-population of 39 cells; 21 cells removed, mouse n= 9). Day 3: mouse(R) decoding = 0.64, null model = 0.50±0.03; location(R) decoding = 0.83, null model = 0.51±0.04; mouse(NS) decoding= 0.61, null model = 0.50±0.03; location(NS) decoding = 0.86, null model = 0.50±0.04. (Pseudo-population of 39 cells; 7 cells removed, mouse n= 4). Day 7: mouse(R) decoding = 0.61, null model = 0.50±0.03; location(R) decoding = 0.73, null model = 0.51±0.04; mouse(NS) decoding= 0.54, null model = 0.51±0.03; location(NS) decoding = 0.73, null model = 0.51±0.03. (Pseudo-population of 39 cells; 15 cells removed, mouse n= 5). **b)** CA2 (purple) mouse-preferring cells are required for mouse identity decoding on days 0 and 3 but not 7. Location-preferring cells are not required for location decoding. Day 0: mouse(R) decoding = 0.82, null model = 0.50±0.03; location(R) decoding = 0.77, null model = 0.49±0.03; mouse(NS) decoding= 0.56, null model = 0.51±0.03; location(NS) decoding = 0.81, null model = 0.50±0.05. (Pseudo-population of 57 cells; 30 cells removed, mouse n= 8). Day 3: mouse(R) decoding = 0.57, null model = 0.50±0.02; location(R) decoding = 0.80, null model = 0.50±0.03; mouse(NS) decoding= 0.53, null model = 0.51±0.03; location(NS) decoding = 0.80, null model = 0.50±0.03. (Pseudo-population of 57 cells; 6 cells removed, mouse n= 5). Day 7: mouse(R) decoding= 0.58, null model = 0.49±0.03; location(R) decoding = 0.89, null model = 0.50±0.05; mouse(NS) decoding= 0.59, null model = 0.50±0.04; location (NS) decoding = 0.89, null model = 0.50±0.03. (Pseudo-population of 57 cells; 4 cells removed, mouse n= 4). Decoding accuracy relative to null model: **p<0.01; ***p<0.001

**Supplemental figure 10.**
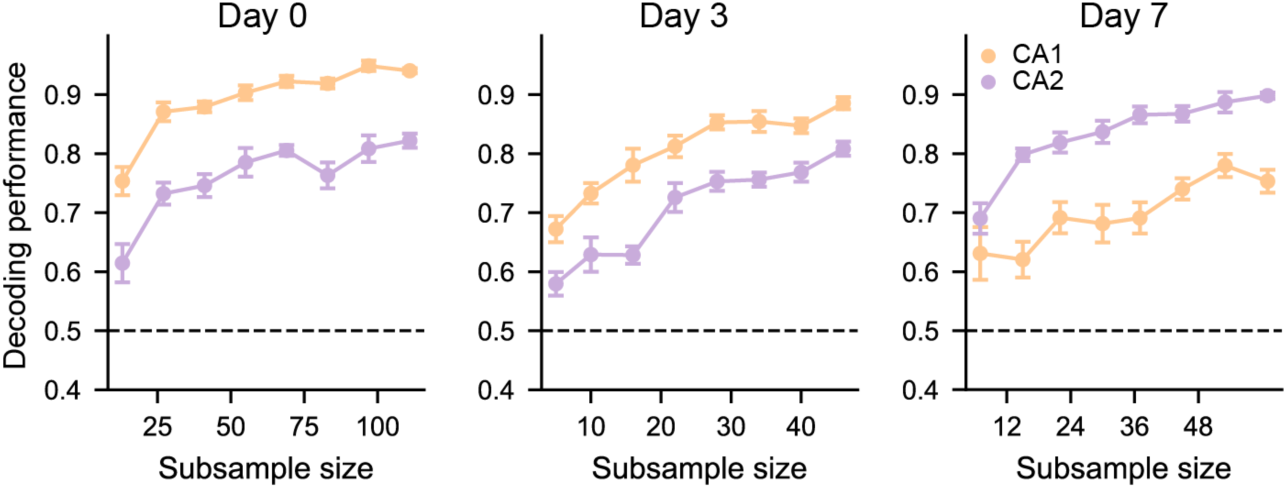
Comparison of location decoding accuracy by CA1 and CA2 pseudo-population subsampling across days. **a)** CA1 exhibits greater location decoding accuracy than CA2 on days 0 and 3; on day 7, CA2 demonstrates greater decoding performance. A linear model examining a region x subsample value interaction revealed that decoding performance increased with subsample value similarly across regions across days (interaction, Type II ANOVA, no significance). (Left) Day 0 Subsamples: 13, 27, 41, 55, 69, 83, 97, 111. (Middle) Day 3 subsamples: 5, 10, 16, 22, 28, 34, 40, 46. (Right) Day 7 subsamples: 7, 15, 22, 30, 37, 45, 53, 61.

